# SARAF represses the mild hypothermia response through the regulation of JUN

**DOI:** 10.64898/2026.09.02.748854

**Authors:** Kijin Jang, Kevin Ostacolo, Valdimar Sveinsson, Michael A. Beer, Kimberley Anderson, Hans Tomas Bjornsson

## Abstract

The mild hypothermia response (MHR) is a conserved mammalian cytoprotective program activated upon exposure to mild hypothermia (32 °C) that contributes to the neuroprotective effects of therapeutic hypothermia following hypoxic injury. Although rapid changes in intracellular calcium occur upon cooling, the mechanisms linking calcium dynamics to the activation of core MHR factors such as SP1 and RBM3 remain incompletely defined. In this study, we used siRNA-mediated knockdown (KD) of candidate regulators in conjunction with novel mild hypothermia indicator (MHI) reporters to identify upstream modulators of MHR-associated transcription. We identify SARAF, a negative regulator of store-operated calcium entry (SOCE), as a repressor of both *SP1* and *RBM3* under normothermic conditions. SARAF depletion is associated with increased intracellular calcium release and enhanced SP1- and RBM3-linked transcriptional outputs. We identify JUN as an important downstream factor mediating SARAF depletion-dependent de-repression of the MHR and demonstrate that it undergoes activation rapidly upon cooling. Finally, SARAF depletion conferred significant cytoprotection against hypoxia-induced early apoptosis. Collectively, these findings establish SARAF as an upstream regulator of MHR-associated transcription and provide a functional link between cold-induced intracellular calcium dynamics and the induction of core MHR effectors.

## Introduction

Human homeostatic mechanisms maintain core temperature within a narrow physiological range (36.2-37.5 °C)^1^. Despite this stringent regulation, the reduction of core temperature to approximately 32 °C is widely applied in clinical medicine as a clinical intervention termed Targeted Temperature Management (TTM), where it is used to minimize neurological damage following neonatal asphyxia^2^ or cardiac arrest^3–4^. Despite its established neuroprotective efficacy, TTM can be accompanied by serious adverse effects^5–7^. At this exact temperature (32 °C), there is activation of an evolutionarily conserved mammalian cytoprotective program termed the mild hypothermia response (MHR)^8^. Exposure to mild hypothermia induces various cellular alterations including a short-lived flux of calcium from the endoplasmic reticulum to the cytoplasm^9,10^ through a calcium channel (ITPR3)^9^ as well as a global attenuation of protein translation, regulated in part, through the activation of eukaryotic elongation factor 2 kinases (eEF2K)^9^. Despite this global decrease in protein production, the transcription and translation of three genes—*SP1*, *CIRBP,* and *RBM3,* respectively encoding specificity protein 1, cold-inducible RNA-binding protein, and RNA-binding motif protein 3—are consistently upregulated at 32 °C^11^. Importantly, this selective induction seems to be central to the neuroprotective effect of mild hypothermia^12–15^.

Current understanding of MHR regulation suggests the presence of two distinct arms of the MHR. First, SP1, a transcription factor (TF) involved in diverse cellular processes^16^, interacts with the promoter region of *CIRBP* during hypothermia. This interaction occurs specifically at a sequence called the Mild Cold Response Element (MCRE)^17^, leading to increased expression of *CIRBP* at 32 °C. Second, RBM3 is known to regulate at least one downstream factor, Reticulon 3 (RTN3), a potential neuroprotective factor validated in two distinct mouse models of neurodegeneration^12–15^. Collectively, these findings suggest that mammalian cells have a regulatory pathway culminating in the coordinated expression and activity of the SP1/CIRBP and RBM3/RTN3 axes; however, the upstream regulatory mechanisms remain incompletely understood.

To begin identifying these upstream regulators, we used a genetic screening approach and created multiple mild hypothermia indicators (MHIs), which translate transcriptional activity of the promoters of three key genes (*SP1*, *CIRBP*, *RBM3*) into discernible fluorescent signals, thereby enabling single-cell quantification of MHR activation^18^. Using one of these MHIs (*SP1*-MHI) in a CRISPR-Cas9 screen, we previously identified a novel repressor of *SP1*, a histone methyltransferase called SMYD5^18^. Recent studies have demonstrated that SMYD5 also trimethylates RPL40, a ribosome subunit component of UBA52, and this modification is necessary for optimal ribosome and translation efficiency^19–21^. Loss of SMYD5 abolishes this methylation, thus impairing the UBA52-dependent ribosomal activity^19–21^. Thus, the degradation of SMYD5 could be another way that hypothermia leads to a general decrease of protein output. However, because SMYD5 selectively regulates SP1, but not RBM3^18^ and given the scarcity of direct links between cold-induced intracellular calcium fluctuations and core MHR factors^11^, we hypothesized the existence of additional, currently unknown distal regulators that converge on both arms of the MHR.

In this study, we have selected candidate regulators from our prior CRISPR-Cas9 screen^18^ and from the literature and screened them using our MHIs. We demonstrate that SARAF (store-operated calcium entry-associated regulatory factor), a regulator of store-operated calcium entry (SOCE), functions as a constitutive repressor of both *SP1* and *RBM3*, thereby playing a role as a repressor of both arms of the MHR. Furthermore, we show that de-repression of the MHR requires the presence of the calcium-responsive TF JUN. Collectively, these findings establish a mechanistic link between cold-induced intracellular calcium dynamics, calcium-responsive TFs, and the regulation of the mammalian MHR.

## Results

### Unbiased screening uncovers SARAF as a novel regulator of both arms of the MHR

Given that mild hypothermic stimuli trigger rapid, early fluctuations in cytosolic calcium^9,10^, we hypothesized the existence of distal, calcium-responsive upstream regulators capable of modulating both arms of the MHR. To identify potential upstream regulators of the MHR, we selected 16 candidates for evaluation using our MHIs and RT-qPCR, including 11 candidate regulators from our prior publication^18^ —ATF2, CABP4, CALHM2, DDX20, HDAC6, HRC, PRKCE, RASD1, SBF1, SYT1, and SARAF— and 5 additional candidate regulators (AKT1, ATF7, BAZ2A, NRF2, and PGC-1α) from the literature^22–24^ or from an unpublished RBM3-Cas9 forward mutagenesis screen. To systematically evaluate their regulatory influence, we performed siRNA-mediated knockdown (KD) of each candidate individually and quantified changes in *SP1*-MHI fluorescence intensity at 37 °C (**Figure 1A**). Independent KD of 6 of these factors (*SARAF*, *HDACC*, *CALHM2*, *CABP4*, *SBF1*, and *PRKCE*) resulted in a significant induction of *SP1*-MHI fluorescence intensity, consistent with a repressive role.

**Figure 1.**
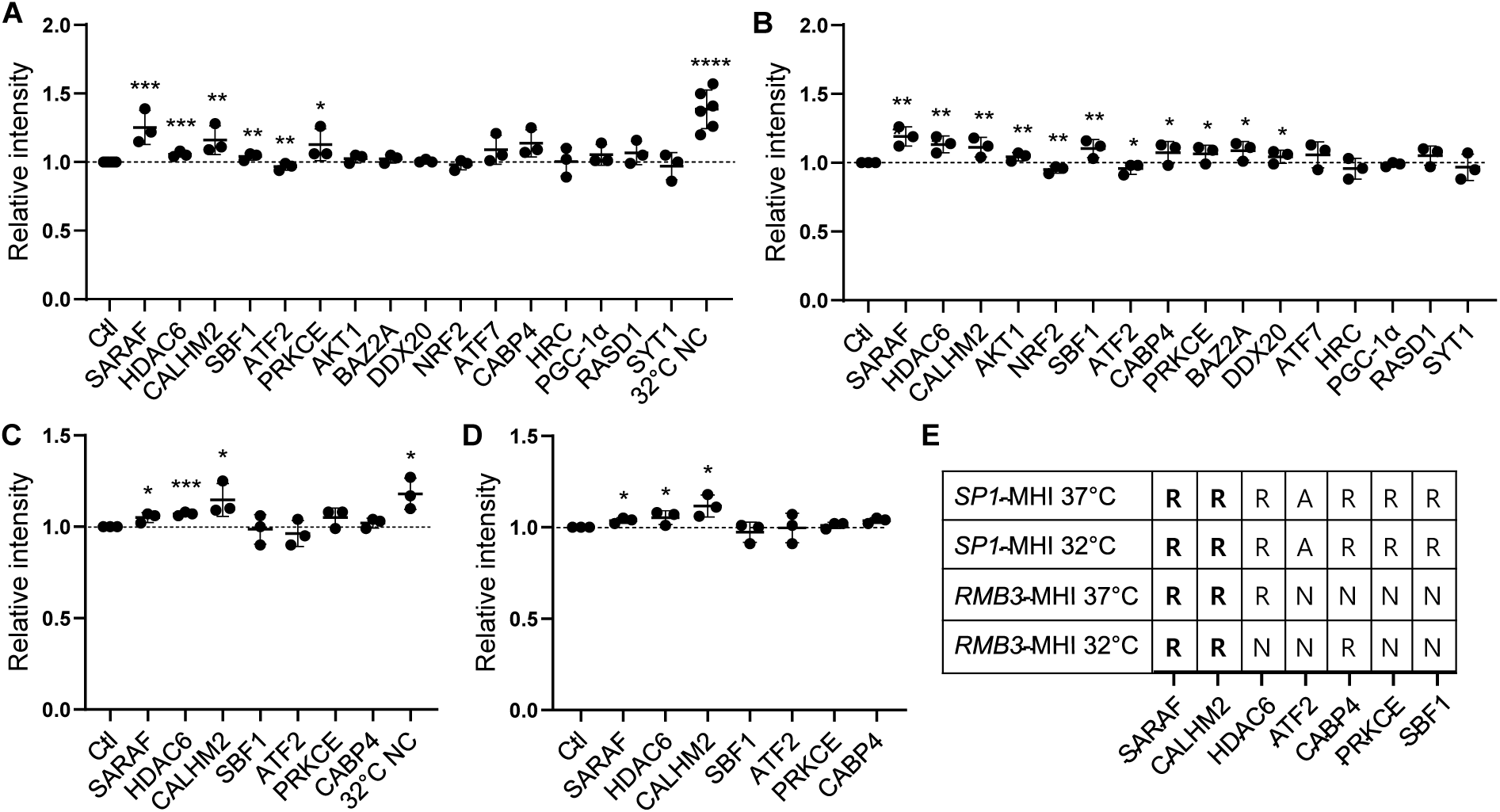
A dual MHI screen to validate candidate upstream regulators of both SP1 and RBM3. (A-B) Flow cytometric profiling of the *SP1*-MHI reporter line, illustrating the effect on fluorescence levels after individual candidate gene silencing at 37 °C (A) and at 32 °C (B). (C-D) Parallel flow cytometric analyses using the *RBM3*-MHI reporter line, demonstrating the regulatory impact of candidate depletion on *RBM3* promoter activity at 37 °C (C) and at 32 °C (D). (E) A summary table of the data from A-D. R, A, and N denote repressors, activators, and non-significant, respectively. Data are displayed as mean ± SD (n = 3–6). Statistical significance was evaluated by a Student’s t-test. Asterisks above individual data points indicate significance in comparison with the controls (NC, 37 °C). *, **, ***, and **** indicate *p* < 0.05, *p* < 0.01, *p* < 0.001, and *p* < 0.0001, respectively.

Conversely, KD of *ATF2* decreased *SP1*-MHI fluorescence intensity, suggesting a role as an activator of *SP1*. Next, we evaluated the impact of these KDs on *SP1*-MHI activity at 32 °C. KD of all seven candidates recapitulated the effects observed at 37 °C (**Figure 1B**). Additionally, KD of *AKT1*, *BAZ2A*, and *DDX20* also induced a significant increase in *SP1*-MHI fluorescence intensity. Again, only ATF2 exhibited activator-like properties consistently, while the others displayed profiles consistent with that of a transcriptional repressor. Based on these results, the seven candidates displaying the most robust and consistent regulatory profiles were selected for additional screening using the *RBM3*-MHI (**Figure 1C-D**). Strikingly, KD of *SARAF* or *CALHM2* increased RBM3-MHI fluorescence intensity at both temperatures. We also observed a significant effect of *CABP4* and *HDAC6* silencing at one of the temperatures, whereas the others did not lead to significant changes. We have summarized the results of our screening in **Figure 1E**, which shows that independent silencing of two genes, *SARAF* and *CALHM2*, consistently showed upregulation in SP1 and RBM3 expression at both temperatures, while SARAF was the only candidate showing statistical significance in *SP1*-MHI fluorescence intensity at both temperatures (*p* < 0.01, one-way ANOVA with a Dunnett’s multiple comparison correction). SARAF (also known as TMEM66), an established negative regulator of SOCE, emerged as the strongest candidate given prior literature on calcium fluxes early upon a hypothermia stimulus^9,10^ and its known role in regulating cytosolic calcium^25^. We thus prioritized SARAF for in-depth mechanistic validation.

### SARAF functions as a repressor of both arms of the MHR

To validate the regulatory role of SARAF in the MHR, *SARAF* was silenced using siRNA and the downstream effects on both arms of the pathway were assessed. The efficiency of *SARAF* KD was confirmed by RT-qPCR at both 37 °C and 32 °C (**Figure 2A**). Under normothermic (37 °C) conditions, *SARAF* depletion resulted in a significant (*p* < 0.01) increase in endogenous *SP1* mRNA levels compared to the non-targeting control (**Figure 2B**). In contrast, no significant difference was observed in *SP1* expression between *SARAF* KD and control cells at 32 °C, indicating that SARAF primarily constrains *SP1* transcription under basal conditions rather than during active hypothermic induction. Similarly, silencing *SARAF* significantly increased *RBM3* mRNA levels at 37 °C, whereas again no significant difference was observed between *SARAF* KD and control cells at 32 °C (**Figure 2C**). These results indicate that SARAF acts as a repressor of both *SP1* and *RBM3* under normothermia. As shown earlier from the results of the screening using MHIs, *SARAF* KD increased MHI fluorescence at 37 °C in both reporter systems, which is consistent with the RT-qPCR findings, indicating the activation of *SP1*- and *RBM3*-linked transcriptional programs upon *SARAF* silencing under normothermia (**Figure 2D–E**). Finally, to determine whether SARAF also regulates SP1 and RBM3 at the protein level, we performed western blot analysis. *SARAF* KD resulted in increased protein levels of both SP1 and RBM3 under normothermic conditions, confirming that SARAF constrains both arms of the MHR not only at the mRNA level but also at the protein level (**Figure 2F–H**). Taken together, RT-qPCR, MHI-based flow cytometry, and western blot analyses consistently demonstrate that SARAF functions as a repressor of both the SP1 and RBM3 arms of the MHR under normothermic conditions.

**Figure 2.**
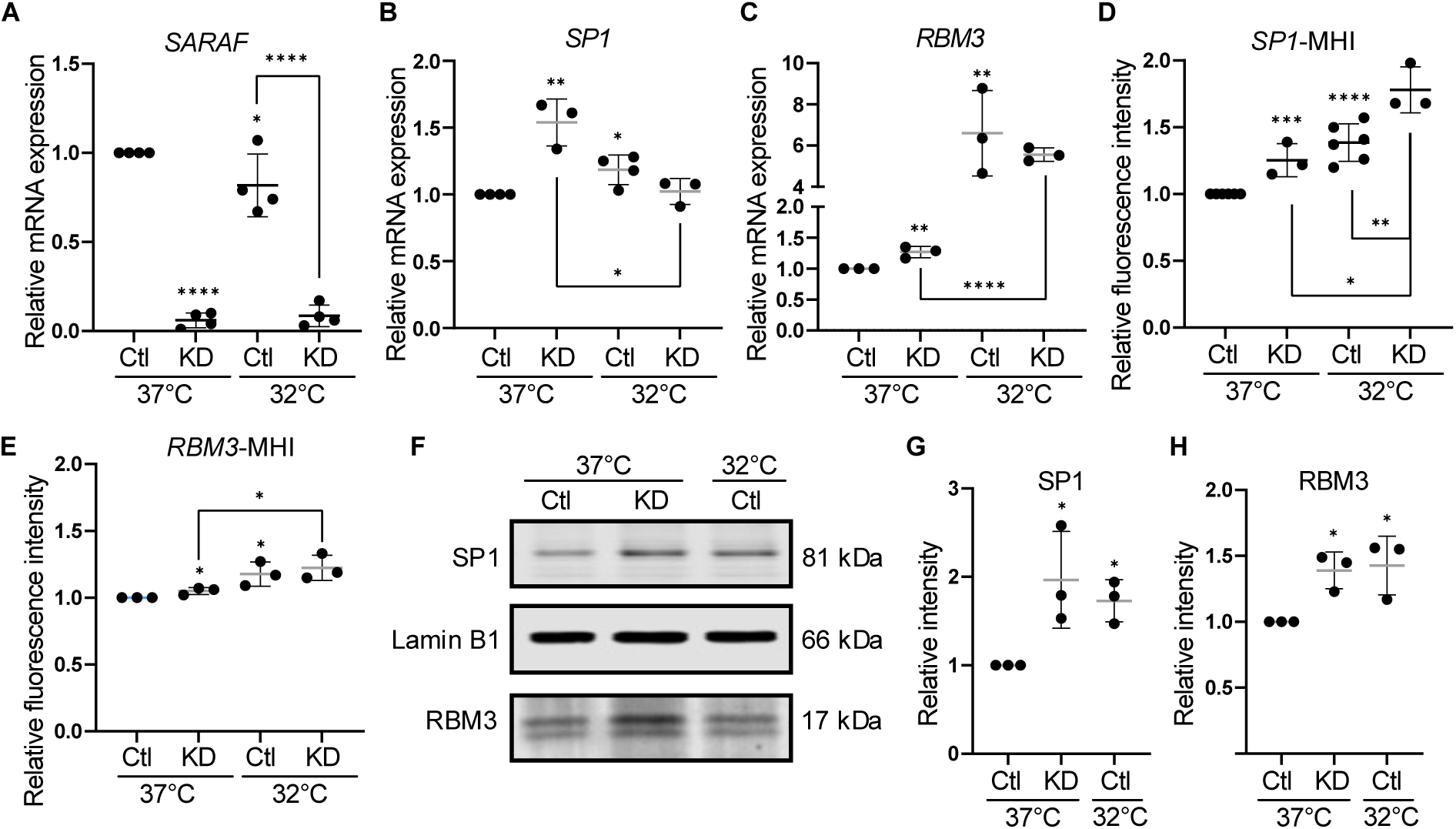
SARAF silencing enhances the endogenous loci of two key MHR effectors under normothermic conditions. (A-C) RT-qPCR validation of siRNA-mediated target silencing, quantifying (A) *SARAF* KD efficiency and the corresponding relative mRNA expression of (B) *SP1* and (C) *RBM3* at 37 °C and at 32 °C. (D–E) Flow cytometric quantification of (D) SP1-MHI and (E) RBM3-MHI reporter line fluorescence intensity at 37 °C and at 32 °C following *SARAF* silencing, confirming elevated promoter activity under normothermic conditions. (F–H) Western blot analysis evaluating downstream translation products, showing (F) representative immunoblots and the corresponding densitometric quantification of (G) SP1 and (H) RBM3 protein abundance at 37 °C. Data are displayed as mean ± SD (n = 3-6). Note that the datasets presented in panels D-E represent a prioritized subset of the broader screen detailed in **Figure 1A-D**. Statistical significance was evaluated via a one-way ANOVA followed by Holm-Šídák’s post-hoc multiple comparisons test for panels A, G, and H and via Student’s t-test for panels B, C, D, and E. Asterisks above individual data points indicate significant differences relative to the controls (NC, 37 °C), while brackets delineate statistically significant differences between specified experimental groups. *, **, ***, and **** denote *p* < 0.05, *p* < 0.01, *p* < 0.001, and *p* < 0.0001, respectively.

### SARAF is a dynamic regulator of intracellular calcium levels

To determine whether SARAF itself is temperature-sensitive, we performed immunofluorescence (IF) staining, revealing a transient decrease in cytoplasmic SARAF intensity during a short-lived (2 h, **Figure 3A-B**) hypothermic stimulus. Notably, this change resolved within 6 hours of the hypothermic stimulus (**Figure 3B**), indicating rapid resolution of the response. Concurrently, spatial granularity analysis revealed a redistribution of the SARAF signal by mild hypothermia over time. At 37 °C, high-order granularity features (e.g., level 12, see methods) were elevated (**Figure 3C, Supplementary Figure 1**), consistent with a diffuse spatial organization. Upon a 2-hour exposure to 32 °C, high-order granularity was markedly reduced (**Figure 3C**), whereas low-order granularity (e.g., level 2) increased (**Supplementary Figure 2**), indicating a shift toward punctate structures. By 6 hours of mild hypothermic exposure, the spatial distribution of SARAF returned to the baseline levels seen in the 37 °C control (**Figure 3C, Supplementary Figures 1-2**). These findings suggest that SARAF undergoes rapid, transient spatial reorganization during the early hypothermic response, potentially reflecting the dynamic remodeling of SOCE-associated signaling complexes.

**Figure 3.**
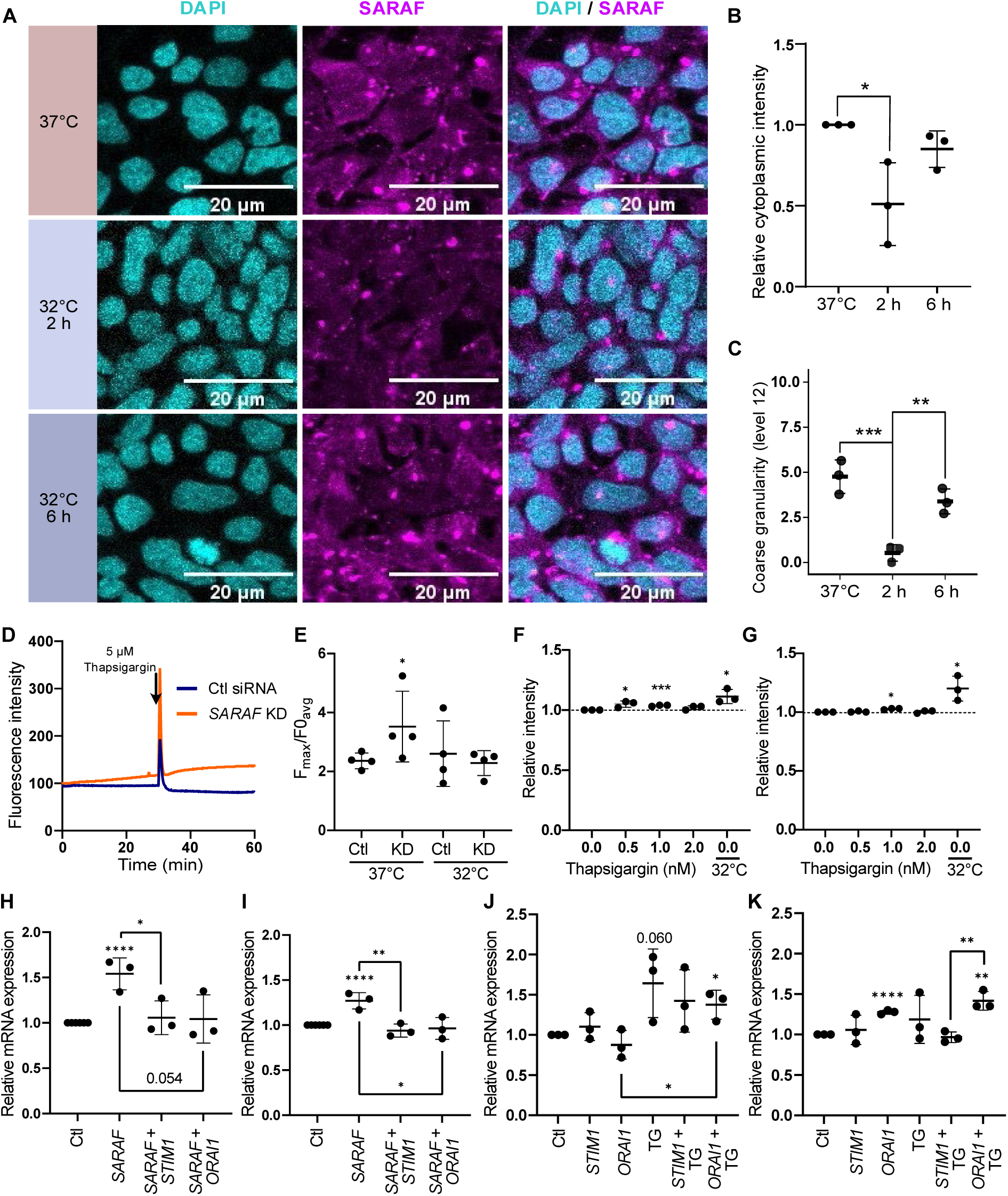
The impact of SARAF silencing on intracellular calcium homeostasis and MHR induction displays a temperature-sensitivity. (A) Representative confocal images of SARAF and DAPI staining in a time-course analysis (0, 2, and 6 hours). (B) Quantitative analysisof cytoplasmic SARAF fluorescence intensity across the individual time points from panel A. (C) Quantification of granularity (i.e., distribution) of SARAF from IF images. (D-E) Fluorometric quantification of dynamic changes in intracellular calcium, using the calcium-sensitive indicator Cal-520. (D) Representative real-time kinetics profiles tracing intracellular calcium fluorescence intensities in HEK293T cells maintained under normothermia with (right) or without (left) *SARAF* KD with TG addition at 30 minutes. (E) Quantification of the TG-induced intracellular calcium release efficiency from multiple traces such as the one shown in D. Flow cytometric profiling of the (F) *SP1*- and (G) *RBM3*-MHIs following a 3-hour TG exposure. (H-I) RT-qPCR analysis evaluating the impact of concurrent KD of *SARAF* and either *STIM1* or *ORAI1* on (H) SP1 or (I) RBM3 mRNA expression. (J-K) RT-qPCR analysis examining the impact of the SOCE regulators KD combined with TG exposure on the mRNA expression of (J) *SP1* and (K) *RBM3*. Data are displayed as mean ± SD (n = 3-4). Note that the data of *SARAF* KD alone and the subset (n = 3-4) of controls at 37 °C presented in **Figures 3H and 3I** are identical to the data of *SARAF* KD and controls at 37 °C described in **Figures 2B and 2C**, respectively. Statistical significance was evaluated via a one-way ANOVA followed by Holm-Šídák’s post-hoc multiple comparisons test for panel B, and by Tukey’s *post*-*hoc* multiple comparisons test for panel C, and via Student’s t-test for panels E-K. Asterisks positioned directly above individual data bars indicate significant variations relative to the controls at 37 °C, while brackets delineate statistically significant differences between specified experimental groups. *, **, ***, and **** denote *p* < 0.05, *p* < 0.01, *p* < 0.001, and *p* < 0.0001, respectively.

Since SARAF undergoes rapid spatial redistribution following mild hypothermic exposure and is an established modulator of cytoplasmic calcium levels^25^, we next investigated whether these early changes alter intracellular calcium dynamics. We used Thapsigargin (TG) both as a standardized ER calcium-release stimulant in calcium measurements and as a pharmacological agent to evaluate whether elevated cytosolic calcium is sufficient to activate MHR transcriptional outputs. Intracellular calcium levels were monitored using the fluorescent calcium indicator Cal-520 in calcium-free HBSS, enabling measurement of cytosolic calcium dynamics in the absence of extracellular calcium influx. Cells were first monitored for 30 minutes to establish baseline fluorescence. Subsequently, TG, a Sarcoplasmic/Endoplasmic Reticulum Calcium ATPase (SERCA) inhibitor that depletes ER calcium stores by inducing calcium release into the cytoplasm, was added and measurements were continued for an additional 30 minutes. Under normothermic conditions (37 °C), *SARAF* KD increased the magnitude of cytosolic calcium release following TG addition compared to control cells (**Figure 3D**). This is similar to what has been seen in prior studies on SARAF depletion in HeLa and Jurkat cells treated with a calcium ionophore, ionomycin^25^. In contrast, at 32 °C, the peak cytosolic calcium amplitude was not notably different between *SARAF* KD and control conditions (**Supplementary Figure 3A**). Quantifying these dynamics across biological replicates (**Figure 3E, Supplementary Figure 3A**) confirmed that *SARAF* KD significantly increased both intracellular calcium release efficiency and peak amplitudes at 37 °C but not at 32 °C (**Figure 3E, Supplementary Figure 3B**). These data suggest that the role of SARAF in calcium regulation is limited to normothermia. Additionally, baseline cytosolic calcium levels were increased under mild hypothermia (**Supplementary Figure 3C**), which is consistent with prior literature^9,10^. Taken together, these results support a model in which SARAF modulates intracellular calcium release dynamics predominantly at 37 °C, consistent with a role as a repressor of the MHR at normothermia.

To test whether experimentally increasing cytosolic calcium is sufficient to activate the MHR in the absence of mild hypothermia, we pharmacologically increased cytosolic calcium using TG. Cells harboring either the *SP1*- or *RBM3*-MHIs were exposed to TG for 3 hours at 37 °C. To minimize cytotoxicity and account for the potential dose-dependent effect of calcium perturbation, three concentrations (0.5, 1, and 2 µM) were tested. TG exposure led to a modest but statistically significant increase in *SP1*-MHI fluorescence at 0.5 and 1 µM (**Figure 3F**). Similarly, in *RBM3*-MHI cells, TG also increased reporter activity, with the strongest and most consistent induction observed at 1 µM (**Figure 3G**). Based on these results, 1 µM TG was selected for subsequent experiments. Collectively, these data demonstrate that acute elevation of cytosolic calcium is sufficient to enhance SP1- and RBM3-linked transcriptional activity under normothermic conditions, indicating that calcium spikes can engage the MHR transcriptional outputs, bypassing the requirement for a hypothermia stimulus.

Given the established role of SARAF as a negative regulator of SOCE^25^, we sought to determine whether this regulatory function contributes to MHR regulation. During SOCE, STIM1 functions as the main ER calcium sensor, which activates the membrane-bound ORAI1 calcium channel in response to depleted ER calcium stores^26^. To evaluate whether SARAF represses the MHR at 37 °C through inhibition of SOCE, we performed combinatorial KD of *SARAF* with SOCE factors *STIM1* or *ORAI1*, and evaluated the impact on transcriptional output of *SP1* (**Figure 3H**) and *RBM3* (**Figure 3I**). This experiment revealed that silencing either *STIM1* or *ORAI1* prevented the de-repression of *SP1* (**Figure 3H**) or *RBM3* (**Figure 3I**) following *SARAF* KD. Conversely, the induction of *SP1* following TG-induced ER calcium release was not dependent on STIM1 or ORAI1, whereas *RBM3* induction was dependent on STIM1 (**Figure 3J-K**). Interestingly, *ORAI1* KD alone induced *RBM3* expression, possibly through a compensatory calcium entry mechanism (**Figure 3J-K**). Collectively, these data indicate that increased cytosolic calcium is sufficient to activate key transcriptional outputs of the MHR, whether achieved through SOCE or through ER calcium release in the absence of SOCE (i.e., upon *ORAI1* or *STIM1* KD).

### SARAF-dependent derepression of SP1 and RBM3 requires JUN

Since SARAF resides in the cytoplasm, whereas the transcriptional activation of *SP1* and *RBM3* occurs in the nucleus, we hypothesized that the effect of SARAF depletion on the MHR is mediated by one or more TFs. To first assess whether mild hypothermia induces broad chromatin remodeling at MHR loci, we performed ATAC-seq at 37 °C and 32 °C at a single time point (7 hours) to identify major changes in chromatin accessibility. We analyzed these data using gkm-SVM^27^ to evaluate enrichment of open chromatin states. Globally, there was a strong overall peak concordance (N=178,404 peaks, R^2^=0.97), indicating the general preservation of chromatin accessibility independent of temperature. However, we observed a modest increase in accessibility at 32 °C compared to at 37 °C (**Supplementary Figure 4A, Figure 4A**). Global motif enrichment analysis within these differentially accessible regions revealed an enrichment of KLF and ZNF motifs in peaks with increased accessibility at 32 °C compared to at 37 °C (**Supplementary Figure 4B**), whereas ATF and MAZ motifs were enriched in peaks showing decreased accessibility (**Supplementary Figure 4B**). Next, to identify the specific TFs driving these alterations, we used CentriMo to identify centrally enriched motifs, which can indicate TFs that bind at peak summits and are therefore more likely to directly contribute to changes in chromatin accessibility^28^. KLF, NFY, AP-1, and ZNF motifs were enriched at the peak centers gaining accessibility (**Figure 4B**), whereas KLF, bHLH, and bZIP motifs were associated with peak centers showing reduced accessibility at 32 °C (**Figure 4B**). The KLF motif can represent a binding site for a large number of SP/KLF-family TFs, including SP1^29^, while the AP-1 motif is recognized by a number of TFs including the immediate-early TF JUN^30^.

**Figure 4.**
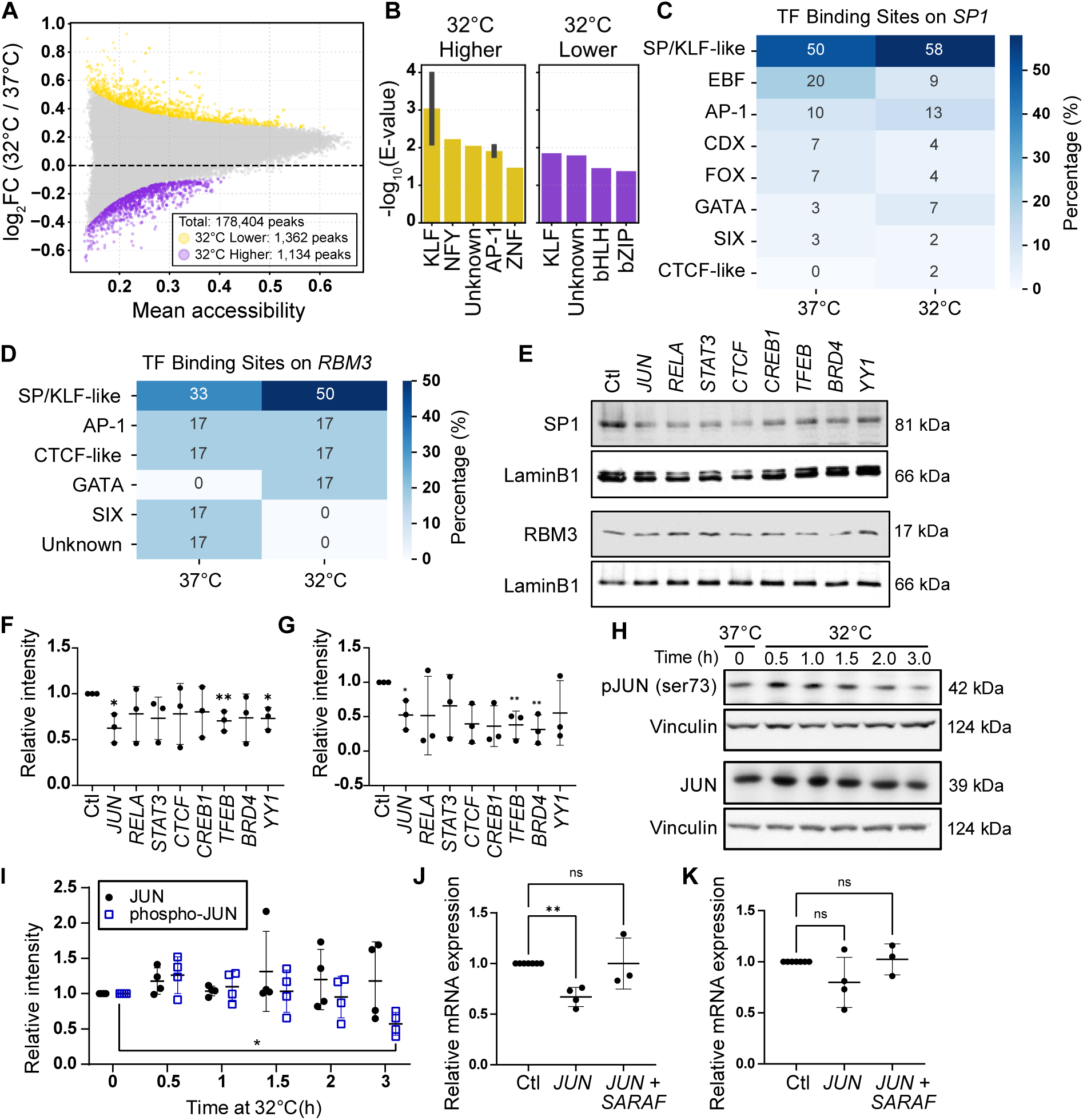
Rapid changes in JUN and its phosphorylation state mediate transcriptional regulation of SARAF. (A) A MA plot of chromatin accessibility comparing 32 °C and 37 °C from an ATAC-Seq experiment. Each point represents an ATAC-seq peak plotted as mean accessibility versus log₂ fold change (32°C/37°C). Yellow indicates sites with increased accessibility at 32 °C compared to 37 °C. Purple indicates increased accessibility at 37 °C compared to 32 °C. (B) TF motif enrichment across accessibility groups identified in A, determined using MEME/FIMO. Values represent the percentage of peaks containing motifs from each TF family in regions with increased or decreased accessibility at 32 °C. Central enrichment of TF motifs within accessible regions, assessed using CentriMo. Bars represent −log₁₀(E-value) for motifs enriched near peak centers, highlighting candidate TFs with potential direct regulatory activity. The color scheme is the same as in A. (C-D) Transcription factor motifs within open chromatin binding motifs at (C) the *SP1* promoter and (D) the *RBM3* promoter. The blue scale indicates the enrichment of individual motifs within open chromatin. (E-G) Western blot analysis demonstrating the expression of (F) SP1 and (G) RBM3 in HEK293T cells in the context of KD of 8 TFs. Representative western blots of SP1 and RBM3 are displayed in panel (E). (H-I) Western blot analysis of JUN and phospho-JUN (Ser73) expression under time-course mild hypothermic conditions. (H) Representative western blots for a time series of JUN and phosphorylated JUN (Ser73). (I) Quantification of multiple replicates of JUN and phosphorylated JUN (Ser73) from panel H. (J-K) RT-qPCR analysis evaluating the impact of concomitant KD of *JUN* and *SARAF* on the mRNA expression of (J) *SP1* and (K) *RBM3*. Data are displayed as mean ± SD (n = 3). Statistical significance was evaluated via a one-way ANOVA followed by Dunnett’s post-hoc multiple comparisons test for panel I and via a Student’s t-test for panels F, G, J, and K. *, **, ***, and **** indicate *p* < 0.05, *p* < 0.01, *p* < 0.001, and *p* < 0.0001.

**Figure 5.**
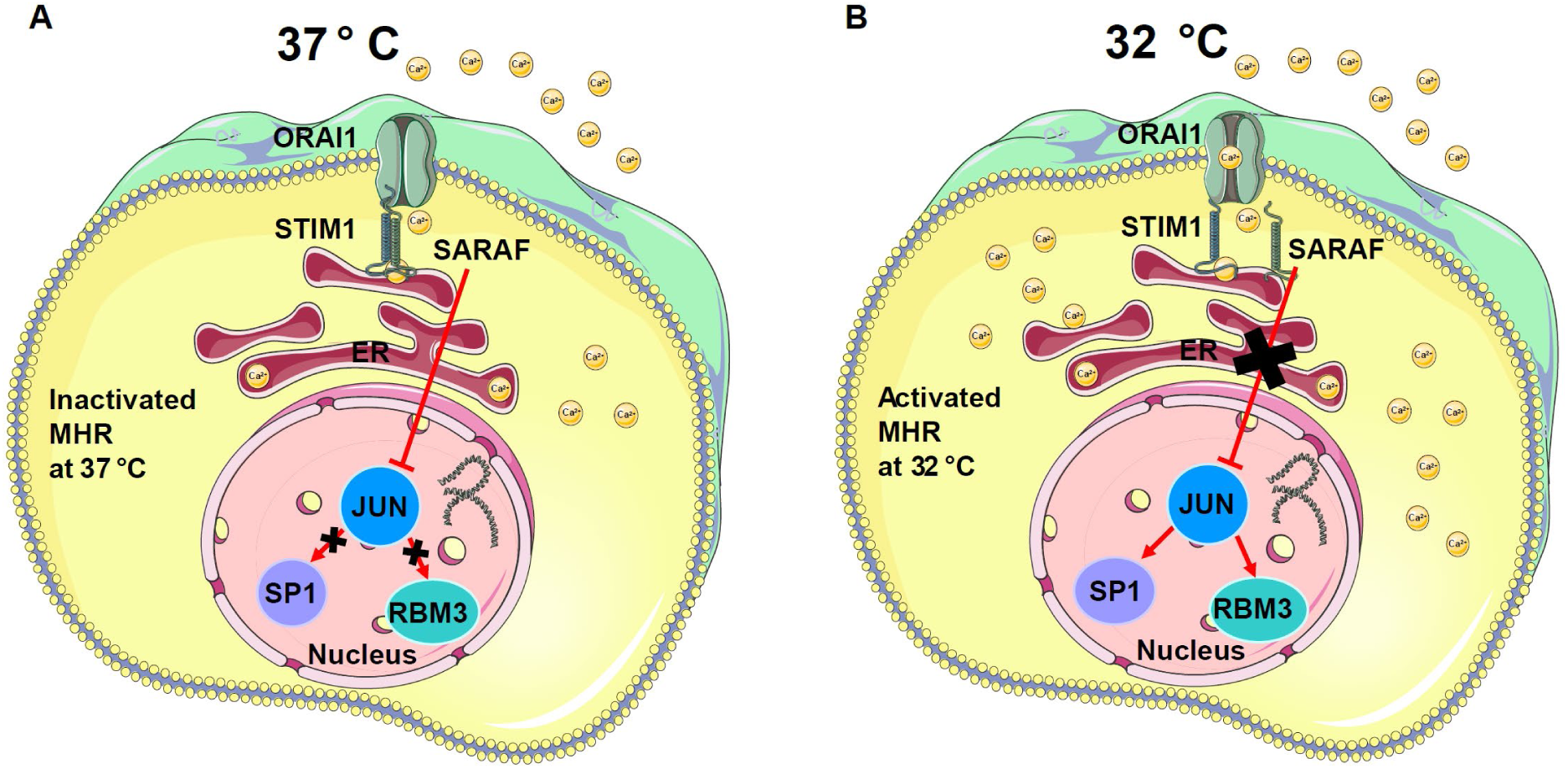
A schematic model of MHR activation regulated by SARAF. (A) Under normothermic conditions SARAF represses the induction of the MHR-induced genes by repressing JUN. When JUN is repressed, it no longer contributes to the activation of SP1 and RBM3. (B) In contrast, in the context of mild hypothermia, SARAF no longer exerts its repressive activity on JUN, which is now able to activate SP1 and RBM3.

To investigate locus-specific regulatory features, we analyzed TF motif composition in ATAC-seq peaks associated with key MHR target genes known to be regulated at the transcriptional level, including *SP1* (**Figure 4C**), *RBM3 (***Figure 4D**) and *CIRBP* (**Supplementary Figure 4C**). Across all loci, SP/KLF-like motifs were highly prevalent, representing over 50% of motifs at 32 °C (**Figure 4C-D and Supplementary Figure 4C**), consistent with the key role of SP1 in the MHR. This was most pronounced at the *CIRBP* locus, where SP/KLF motifs accounted for > 90% of motifs, suggesting strong and potentially constitutive regulatory input, in line with prior reports of SP1-dependent *CIRBP* regulation^17^ (**Supplementary Figure 4C**). AP-1 motifs were also detected across key MHR loci, with modest enrichment at 32 °C at the *SP1* locus, indicating a potential contribution to the broader hypothermic response, consistent with prior literature^31^, while no difference in AP-1 accessibility was noted at *RBM3* (**Figure 4D**). Notably, GATA motifs were specifically observed at 32 °C at the *RBM3* locus (**Figure 4D**). In contrast, EBF and SIX motifs were more prominent at 37 °C and were reduced or absent at 32 °C across multiple loci (**Figure 4C-D**).

We next examined the most extreme temperature-dependent changes in accessibility, defined by the 0.01 and 0.99 quantiles (∼1,800 peaks in each direction; **Supplementary Table 1**). Among regions with higher accessibility at 32 °C, 84% were promoter-associated, compared with only 16% of regions with higher accessibility at 37 °C. Conversely, intronic regions accounted for 54% of peaks with higher accessibility at 37 °C, compared with 9% at 32 °C. Thus, the strongest temperature-dependent changes in chromatin accessibility differed markedly in their genomic distribution.

Taken together, these findings indicate temperature-dependent differences in both the genomic distribution of accessible regions and their associated TF motifs, characterized by prominent SP/KLF motif representation and a context-specific enrichment of AP-1 motifs under mild hypothermia. Given the rapid calcium alterations induced by mild hypothermia and SARAF perturbation, we next sought to identify calcium-responsive TFs capable of linking these early calcium signaling events to the transcriptional activation of the MHR. To achieve this, we focused on TFs known to be calcium-responsive and/or previously reported to bind to *SP1* and *RBM3* promoters. Guided by our ATAC-seq data, we selected nine candidate TFs (JUN, CTCF, CREB1, RELA, STAT3, BRD4, YY1, TFEB, and NRF2). These candidates include representatives of bZIP factors (JUN, CREB1, NRF2), consistent with enrichment of AP-1/ATF motifs, as well as zinc finger TFs (CTCF, YY1). TFs were selected based on literature supporting their calcium-dependence^32,33^ and evidence of binding at key MHR factors in ChIP-Atlas^34^ (a public ChIP-seq aggregation database; **Supplementary Figure 5**), and were screened using *SP1*- and *RBM3*-MHIs (**Supplementary Figure 6A-D**) at both temperatures. The individual knockdown of *JUN*, *CTCF*, *RELA*, and *NRF2* resulted in significant changes in reporter fluorescence under at least one temperature. For the RT-qPCR analysis examining the endogenous *SP1* and *RBM3* expression following TF KD (**Supplementary Figure 6E-H**), the effects were generally subtle, with only *CREB* KD inducing a statistically significant reduction in the mRNA level of SP1 under mild hypothermic conditions. Given that many TFs are regulated at the protein level^35^, we next assessed SP1 and RBM3 protein abundance following individual KD of eight candidate TFs, excluding NRF2, which appears to have an opposite role (i.e., an activator of the MHR) to the other candidates, at 32 °C. Under these conditions, western blot analysis revealed that independent silencing of *TFEB* and *JUN* reduced both SP1 and RBM3 expression, whereas *YY1* KD reduced SP1, and *BRD4* KD significantly reduced RBM3 protein levels (**Figure 4E-G**).

Among these candidates, JUN was of particular interest given the western blot analysis results showing the impact of its KD on both SP1 and RBM3, its established calcium sensitivity, and its enriched binding motif at MHR loci under hypothermia, as well as the previously described AP-1 enrichment both globally and at key MHR loci (**Figure 4B-D**). We, therefore, examined the role of JUN within the context of SARAF-dependent regulation.

To explore the temperature-dependent kinetics of JUN, we initially examined nuclear JUN abundance following mild hypothermic (32 °C) exposure using immunofluorescence. Nuclear JUN intensity showed only modest changes over the 2- and 6-hour time points (**Supplementary Figure 7**), prompting us to examine earlier time points and JUN phosphorylation as a more direct measure of activation. Because immediate-early TFs frequently undergo rapid, transient activation kinetics^36^ following calcium signaling, we performed a time-course western blot analysis in 30-minute increments after mild hypothermic exposure, measuring total JUN and phosphorylated JUN (Ser73) (**Figure 4H**). Quantitative analysis across multiple biological replicates (**Figure 4I**) revealed a trend toward an increase in both total JUN and phosphorylated JUN at the initial 30-minute time point. After 3 h, phosphorylated JUN was significantly below the baseline, confirming a rapid, short-lived activation profile.

To determine whether JUN is required for the transcriptional consequences of SARAF depletion, we performed dual-KD experiments. While previously, we observed KD of *SARAF* to induce *SP1* and *RBM3* expression (**Figure 2B-H**), concurrent KD of *SARAF* and *JUN* abrogated the upregulation of *SP1* (**Figure 4J**) and *RBM3* (**Figure 4K**) mRNA expression. Collectively, these data indicate that JUN acts as a calcium-responsive TF that responds to mild hypothermia and is required for SARAF-mediated derepression of *SP1* and *RBM3*. Importantly, JUN phosphorylation was observed within 30 minutes of mild hypothermic exposure, preceding the downstream induction of canonical MHR effectors and placing JUN activation temporally between the early calcium response and subsequent MHR activation.

### A model of how SARAF integrates with the previously defined factors of the MHR

Finally, we present a model of the molecular architecture of MHR suppression under normothermic conditions through crosstalk between the negative SOCE regulator SARAF and a transcription factor JUN. Under basal conditions, SARAF likely regulates the STIM1-ORAI1 complex to exert an inhibitory effect on the transcription factor JUN. This suppression prevents the downstream activation of SP1 and the cold-shock protein RBM3, effectively maintaining the MHR in an “off” state.

## Discussion

Our prior CRISPR-Cas9 MHI screen^18^ identified SMYD5 as a selective repressor of *SP1*, exhibiting no regulatory influence over *RBM3*^18^. In contrast, SARAF depletion derepresses both *SP1* and *RBM3*, thereby identifying SARAF as an upstream regulator capable of simultaneously engaging both canonical arms of the MHR. This distinction suggests that SOCE may offer a strategic target for modulating MHR-associated transcriptional programs. However, it remains unclear whether the coordinated activation of both regulatory arms confers greater cytoprotection than the activation of individual effectors such as RBM3, which currently has the most robust evidence for cytoprotective efficacy^12–15, 37–38^.

Our mechanistic analyses position JUN as a necessary mediator linking SARAF-dependent calcium dynamics to the transcriptional activation of SP1 and RBM3. The activation kinetics of JUN are early, rapid, and transient, effectively bridging immediate cold-induced calcium fluctuations to the downstream activation of canonical MHR effectors. Dual-KD experiments demonstrated that the derepressive effect of *SARAF* depletion is abrogated in the absence of JUN, indicating that JUN is required for this pathway cascade. However, it is notable that knockdown of TFEB, a well-characterized stress-induced TF^39^, also demonstrated measurable effects on both SP1 and RBM3 protein levels in our candidate validation experiments. Given that calcium signaling is known to influence multiple TFs in parallel^40^, it is plausible that MHR activation reflects coordinated input from a network of several calcium-responsive regulators rather than an exclusive dependence on a single factor. While our data establish a requirement for JUN in de-repression of the MHR following *SARAF* KD, additional factors such as TFEB may contribute to shaping the magnitude or context-specific dynamics of the response.

An important mechanistic question is whether these observed phenotypic effects reflect the specific regulation of SOCE or the broader modulation of intracellular calcium homeostasis. As SARAF is a negative regulator of STIM1–ORAI1-mediated calcium entry^41^, the observation that the concurrent KD of either *STIM1* or *ORAI1* attenuated the transcriptional consequences of *SARAF* depletion supports the involvement of canonical SOCE components. Notably, STIM1 has been reported to exhibit temperature-dependent activation^42^, which may be relevant given the thermal sensitivity of the MHR. Together, these findings suggest that SOCE machinery participates in MHR regulation, while the critical determinant may ultimately be the integrated intracellular calcium dynamics rather than any single channel component in isolation.

Although SARAF emerged as the most robustly validated regulator within our dual-reporter framework, multiple additional candidates showed reproducible, albeit generally subtle, effects predominantly on SP1 and most consistent with repressive activity. These observations may indicate that the regulation of the MHR relies on the integrated actions of many distributed regulators rather than a single master hub. Notably, a number of candidates originating from our prior genetic screen suggest that additional regulators of the MHR remain to be uncovered. Among these, knockdown of CALHM2, an ATP-channel protein^43^, also showed consistent regulation across both reporters and temperature conditions and may represent a particularly promising candidate for future mechanistic investigation. The predominance of repressive hits likely reflects, at least in part, the design of fluorescence-based CRISPR screening, which is inherently more sensitive to loss-of-repressor events that increase reporter signal than to identification of activators. Finally, it is worth noting the alternative possibility that disruption of multiple cellular regulators may elicit stress responses that partially converge on SP1-dependent transcription^44^, thereby mimicking aspects of MHR activation.

Collectively, our findings identify SARAF as a distal regulator of the MHR and JUN as an important downstream mediator, linking intracellular calcium regulation and the activation of core mild hypothermia-responsive cytoprotective transcriptional programs.

## Materials and methods

### Statistical considerations

All data of this study are displayed as the mean ± standard deviation (SD) derived from at least three independent biological replicates. Visualization and statistical analyses were performed using GraphPad Prism 10 software. Comparisons between different groups were evaluated by Student’s t-test, or one-way ANOVA followed by Holm-Šídák’s multiple comparisons test. Statistical significance was defined as a *p*-value under 0.05. Calculated *p*-values are graphically represented by asterisks corresponding to the following tiers of statistical significance: *: p < 0.05, **: p < 0.01, ***: p < 0.001, and ****: p < 0.0001.

### Cell culture

Human embryonic kidney (HEK293 and HEK293T) cells and *SP1*-MHI and *RBM3*-MHI reporter cell lines were cultured in Dulbecco’s Modified Eagle Medium (DMEM)/F-12 supplemented with 10% fetal bovine serum (FBS) and maintained at 37 °C in a humidified incubator containing 5% CO_2_. Neomycin (G418/Geneticin) (Santa Cruz, sc-29065A) was used for antibiotic selection of SP1- and RBM3-MHI HEK293T cells previously described^18^.

### TG exposure

To evaluate the functional impact of elevated intracellular calcium on inducing the MHR, HEK293T cells harboring *SP1*-MHI and *RBM3*-MHI were seeded at 4 x 10^5^ cells per well in 12-well plates. Upon achieving approximately 70% confluency, positive control plates were transferred to a humidified incubator maintained at 32 °C and incubated for 16 hours. Concurrently, experimental groups were subjected to 0.5, 1, and 2 µM TG and incubated for 3 hours, while DMSO was added to the control cells. Following the 3-hour incubation, cells were harvested and processed as detailed in the method of flow cytometry. Based on dose-response profiles, a final concentration of 1 µM TG was selected to systematically validate the potential of the SOCE regulators, including STIM1 and ORAI1, on inducing the MHR through increased intracellular calcium. To determine the requirement of the core SOCE regulators, 1 µM TG was added to HEK293T cells, which were incubated for 24-48 hours after being reverse-transfected with siRNAs (*STIM1*, *ORAI1*), upon reaching 70-80% confluency. Cells were incubated for 3 hours following the TG addition, then harvested in TRIzol and processed as detailed in the method of RT-qPCR.

### Reverse transfection using siRNAs

To validate the roles of *SARAF*, *JUN*, *STIM1*, and *ORAI1* in the MHR, silencing these genes was conducted using predesigned dsiRNAs targeting SARAF (IDT, hs.Ri.SARAF.13.2), JUN (IDT, hs.Ri.JUN.13.3), STIM1 (IDT, hs.Ri.STIM1.13.1), and ORAI1 (IDT, hs.Ri.ORAI1.13.1), negative control dsiRNA (IDT, 51-01-14-04), and Lipofectamine 3000 (Thermo Fisher, L3000015), following the manufacturer’s instructions of Lipofectamine 3000. The list of siRNAs used in this study is described in **Supplementary Table 2**.

### Real-time quantitative reverse transcription polymerase chain reaction (RT-qPCR) analysis

Total RNA was isolated from cells maintained at 32 °C or 37 °C using the Direct-zol RNA Microprep kit (Zymo Research), following the manufacturer’s instructions. The concentration and purity of the isolated RNA were measured using a NanoDrop One Microvolume UV-Vis Spectrophotometer (Thermo Fisher Scientific). Subsequently, cDNA synthesis was performed with the High-Capacity cDNA Reverse Transcription Kit (Applied Biosystems) on a MiniAmp Thermal Cycler (Applied Biosystems). qPCR was performed using Luna Universal qPCR Master Mix on the CFX384 Real-Time PCR Detection System (Bio-Rad). Every RT-qPCR assay in this study was conducted in technical triplicate, with samples loaded in 384-well plates. To ensure analytical precision, any replicate exhibiting a cycle threshold (Ct) deviation exceeding 0.5 was excluded from the calculation of the mean Ct values. Detailed sequences for the primers employed in this study are listed in **Supplementary Table 3**.

### Cell lysis

Cells were rinsed with ice-cold phosphate-buffered saline (PBS) and subsequently lysed on ice in a protein extraction buffer comprising RIPA buffer (50 mM Tris-HCl, 1% NP-40, 150 mM NaCl, 0.1% SDS, 2 mM EDTA, 1% sodium deoxycholate), supplemented with phosphatase inhibitor (5870S, Cell Signaling) and protease inhibitor (11836170001, Roche) cocktails. The lysates were incubated on ice for 30 minutes and centrifuged at 16,000 x g for 30 minutes at 4 °C. Following centrifugation, the supernatants were collected and protein concentrations were determined using the Pierce BCA Assay Kit (Thermo Fisher). The supernatants were mixed with 4X loading buffer (Li-Cor) and denatured at 95 °C for 5 minutes.

For analysis of JUN and JUN-phospho-serine 73 protein levels, cells were cultured in 35 mm round plates and grown at 37°C. The plates were then transferred to an incubator set to 32°C for the indicated period of time (30 min, 1 h, 90 min, 2 h, 3 h, 16 h). The cells were harvested following incubation at 32 °C for the indicated time periods. The cells were briefly washed in PBS, then lysed in 100 µL of protein sample buffer (60 mM Tris-HCl, pH 6.8, 2% SDS, 10% glycerol, 0.01% bromophenol blue and 1.25% β-mercaptoethanol). The cell lysate was transferred to a 1.5 mL tube and heated at 95 °C for 5 minutes, followed by cooling on ice for 5 minutes. The samples were then treated with 0.5 µL GENIUS™ Nuclease (Santa Cruz Biotechnology) and incubated at room temperature for 10 minutes, before storage at -20°C.

### Western blot

Equal amounts of protein samples (20-30 µg) were loaded onto gels, separated via SDS-PAGE, and electrotransferred onto polyvinylidene fluoride (PVDF) membranes. The membranes were blocked in 5% bovine serum albumin (BSA) in Tris-buffered saline with 0.1% Tween-20 (TBST) for 1 hour. Membranes were subsequently incubated overnight at 4 °C with the following primary antibodies: rabbit polyclonal anti-SARAF (1:500, Invitrogen, PA5-24237), rabbit polyclonal anti-RBM3 (1:1,000, Proteintech, 14363-1-AP), rabbit polyclonal anti-SP1 (1:1,000, Proteintech, 21962-1-AP), mouse monoclonal anti-Lamin B1 (1:5,000, Proteintech, 66095-1-Ig), rabbit polyclonal anti-JUN (1:5,000, Proteintech, 24909-1-AP), rabbit monoclonal anti-phospho-JUN-Ser73 (1:5000, Proteintech, 80086-1-RR), mouse monoclonal Anti-Vinculin (1:3,000, Sigma, V9131) diluted in TBST with 5% BSA. Following repeated washes in TBST, membranes were incubated for 90 minutes at room temperature with IRDye 800CW donkey anti-rabbit IgG and 680RD donkey anti-mouse secondary antibodies. After three washes with TBST, the protein bands were visualized using the Odyssey CLx Infrared Imaging System. For the JUN and phospho-JUN-Ser73 blots, the membranes were incubated for 60 minutes at room temperature with Goat Anti-Rabbit IgG HCL HRP (1:15,000, Abcam, ab6721), then washed 3x with TBST and visualized using Clarity Max Western ECL Substrate (Bio-Rad) and imaged on the ChemiDoc XRS+ Molecular Imager (Bio-Rad). Bands from all experiments were quantified with ImageJ software.

### Immunofluorescence

HEK293 cells were seeded at a density of 1 x 10^4^ cells per well in 8-well chamber slides (354118, Falcon). Upon reaching approximately 70% confluency, two slides were transferred to 32 °C and incubated for 2 or 6 hours, respectively. After the 2- and 6-hour incubation periods, the culture medium was removed, and the cells were rinsed with PBS. Fixation was performed using 100 µL 4% paraformaldehyde (PFA) for 15 minutes at room temperature, followed by subsequent washes with PBS. Cellular membranes were permeabilized with 0.3% Triton X-100 in PBS (PBST) for 3 minutes at room temperature, followed by blocking with blocking buffer composed of 0.1% PBST and 5% NGS for 1 hour at room temperature. Primary antibodies including rabbit polyclonal anti-SARAF 1:200, Invitrogen, PA5-31588; rabbit polyclonal anti-JUN, 1:200, Proteintech, 24909-1-AP, rabbit anti-STIM1 1:200, Invitrogen, PA5-20371, rabbit anti-ORAI1 1:200, Invitrogen, PA5-109270) were applied and incubated at 4 °C for 24 hours. After the 24-hour incubation, cells were rinsed with 0.1% PBST and incubated with the goat anti-rabbit Alexa Fluor 647 (A-21244, Thermo Fisher) secondary antibody (1:1,000), in blocking buffer at room temperature in the dark for 1 hour. Then, cells were washed with 0.1% PBST followed by PBS. Following additional washes with 0.1% PBST and with PBS, Fluoromount-G Mounting Medium with DAPI (Invitrogen) was applied to each well, which was then sealed with coverslips. The slides were kept at room temperature overnight and subsequently stored at 4 °C. High-resolution imaging was conducted using an Evident FV4000 confocal laser scanning microscope (Olympus).

### Image analysis

Image analysis and quantification were performed using FIJI/ImageJ and quantified by CellProfiler, respectively. Raw .oir immunofluorescence images were first processed in Fiji/ImageJ using a custom macro. Images were imported using the Bio-Formats Importer and maximum intensity projections were generated from z-stacks. Individual fluorescence channels were separated, pseudocoloured and contrast-adjusted using consistent minimum and maximum intensity settings across all samples. Processed channel images were then exported for downstream quantitative analysis.

Quantitative image analysis was performed using CellProfiler version 4.2.6. Processed PNG images were imported and metadata, including temperature, time point and replicate number, were extracted from the filenames. DAPI and SARAF channels were identified and images were grouped according to temperature, time point and replicate. Nuclei were segmented from the DAPI channel using the IdentifyPrimaryObjects module with global Minimum Cross-Entropy thresholding, while SARAF-positive objects were segmented from the SARAF channel using Otsu thresholding. SARAF objects were related to DAPI-positive nuclei using the RelateObjects module. Fluorescence intensity, area occupied, granularity and radial intensity distribution measurements were extracted, and SARAF area was normalized to DAPI area. Measurements were exported as CSV files for downstream analysis.

### Flow cytometry

Cells were rinsed with PBS and harvested using trypsin. After stopping the trypsinization activity by adding medium, cell suspension was subsequently centrifuged at 500 x g for 5 minutes. The cell pellets were rinsed with PBS and centrifuged under identical conditions. For analysis, pellets were resuspended in 500 µL fluorescence-activated cell sorting (FACS) buffer (PBS supplemented with 5% FBS and 0.2% EDTA). Samples were transferred to FACS tubes for acquisition on a BD FACSymphony A3. For the validation of JUN and a panel of candidate regulators (*RELA*, *CTCF*, *CREB1*, *TFEB*, *BRD4*, *YY1*, *STAT3*, *NRF2*), 100 µL of samples were processed in 96-well plates and measured using the BD FACSymphony A3.

### Intracellular calcium measurement

To evaluate the dynamics of intracellular calcium following SARAF silencing at two different temperatures, at 37 °C and 32 °C, cells were loaded with 8 µM Cal-520 (Abcam, ab171868), and 0.02% Pluronic F-127 in calcium-free media incubated for 2 hours. The Cal-520 dye working solution was replaced with fresh calcium-free HBSS to minimize background fluorescence from extracellular fluorophores. Baseline and stimulated intracellular calcium concentrations were measured on the GloMAX detection system at excitation/emission wavelengths of 490/525 nm at 37 °C or 32 °C. Fluorescence intensity was recorded at 0.1 minute intervals for an initial duration of 30 minutes. 5 µM TG (Abcam, ab147487) was added at the 30-minute time point. The measurement was continued for an additional 30 minutes to capture the temporal profile of calcium release.

### ATAC-seq library construction and sequencing

A total of 45,000 HEK293 cells were harvested after 7 hours of incubation at 32°C. Cell lysis, nuclei preparation, and tagmentation were performed using the Zymo-Seq ATAC Library Kit (Zymo Research, D5458) according to the manufacturer’s instructions. The tagmentation step was modified by extending the incubation time at 37°C from 30 min to 60 min. DNA concentration was quantified using the Qubit® dsDNA HS Assay Kit (Thermo Fisher Scientific, Q33231). Libraries were purified using AMPure XP beads (Beckman Coulter, A63881), and library quality, fragment size distribution, and purity were assessed using an Agilent Bioanalyzer with the Agilent High Sensitivity DNA Kit (Agilent Technologies, 5067-4626). Libraries were sequenced on an Illumina NovaSeq X using 150 bp paired-end sequencing, generating 228.5-606.7 million raw read pairs per sample. Quality was assessed using FastQC^45^. Reads were trimmed to 50 bp using BBduk from BBtools to remove adapters, aligned to hg38 with Bowtie2^46^, and reads were de-duplicated. Peaks were called using MACS2^47^, pooling biological replicates for each condition. Peak calling was performed with the human genome size setting (-g hs), no model building (--nomodel), a significance threshold of p = 1 × 10⁻⁹, and signal tracks generated with --SPMR and -B. We then used gkm-SVM to classify condition-specific accessible regions and identify DNA sequence features predictive of regulatory activity^27^.

### ATAC-seq signal comparison and peak classification

To identify differentially accessible peaks between 32 °C and 37 °C, we modeled the global relationship between ATAC-seq signals using linear regression (LinearRegression, scikit-learn). For each peak, we computed the residual between observed and predicted accessibility at 32 °C. Peaks were classified as differentially accessible if their residual exceeded ±2.5 standard deviations from the mean residual distribution. To ensure biological relevance, we further required a minimum absolute difference in accessibility between conditions (|Δ accessibility| > 0.025). Peaks with positive residuals were classified as having higher accessibility at 32 °C, while those with negative residuals were classified as having lower accessibility. For secondary analysis of the most extreme temperature-dependent changes, peaks in the upper and lower 1% of the distribution of accessibility differences between 32 °C and 37 °C were selected and annotated by genomic feature using ChIPseeker^49^.

### ATAC-Seq TF binding at MHI targets

To investigate TF binding potential at key MHI target loci (SP1, SMYD5, CIRBP, and RBM3), we analyzed ATAC-seq peaks overlapping these genes for motif content. Peak sequences were scanned for known TF binding motifs using motif discovery/scanning tools (MEME/FIMO)^48^. For each gene, the proportion of peaks containing motifs from major TF families was calculated separately for 37 °C and 32 °C conditions. TF families were grouped based on motif similarity, SP and KLF motifs were grouped due to their shared recognition of GC-rich binding sequences (GC-box motifs), which are not distinguishable at the sequence level. Results were expressed as the percentage of peaks containing at least one motif from each family.

## Supplementary Tables

Supplementary Table 1. A list of most extreme ATAC-Seq peaks (see separate file)

**Supplementary Table 2.**
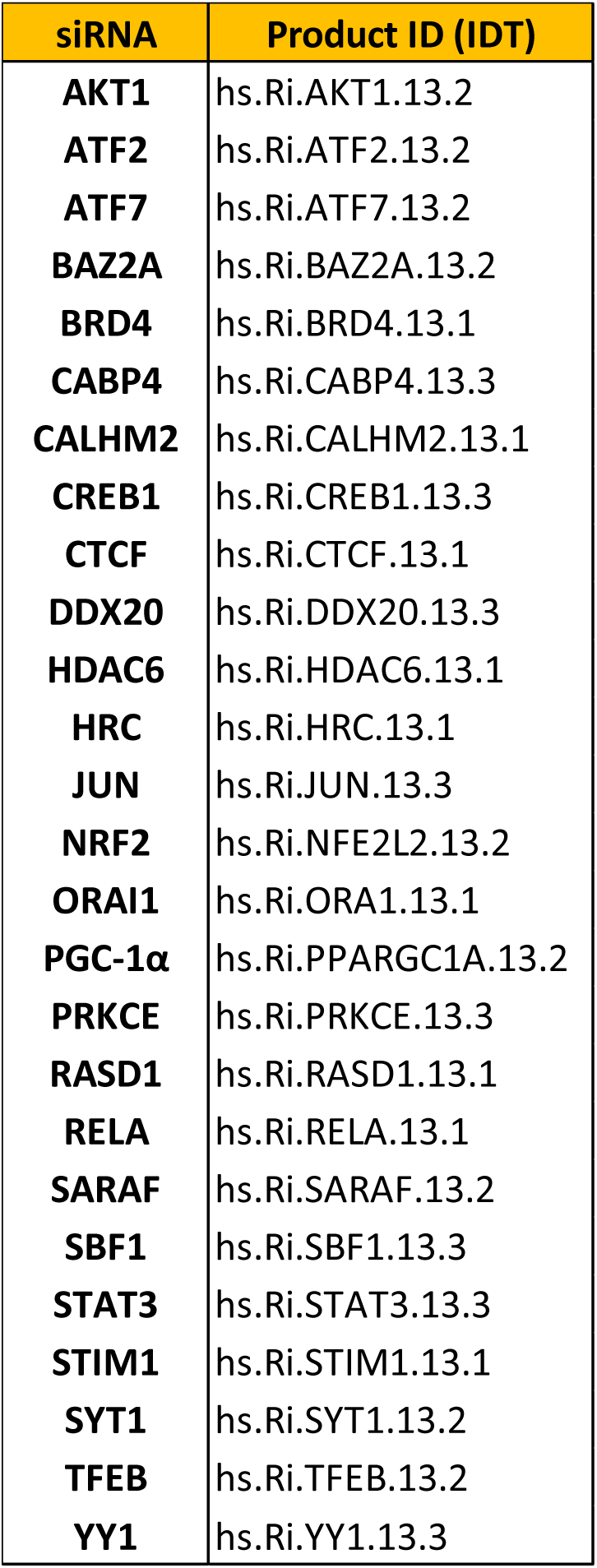
The list of siRNAs used for KD during screening.

**Supplementary Table 3.**
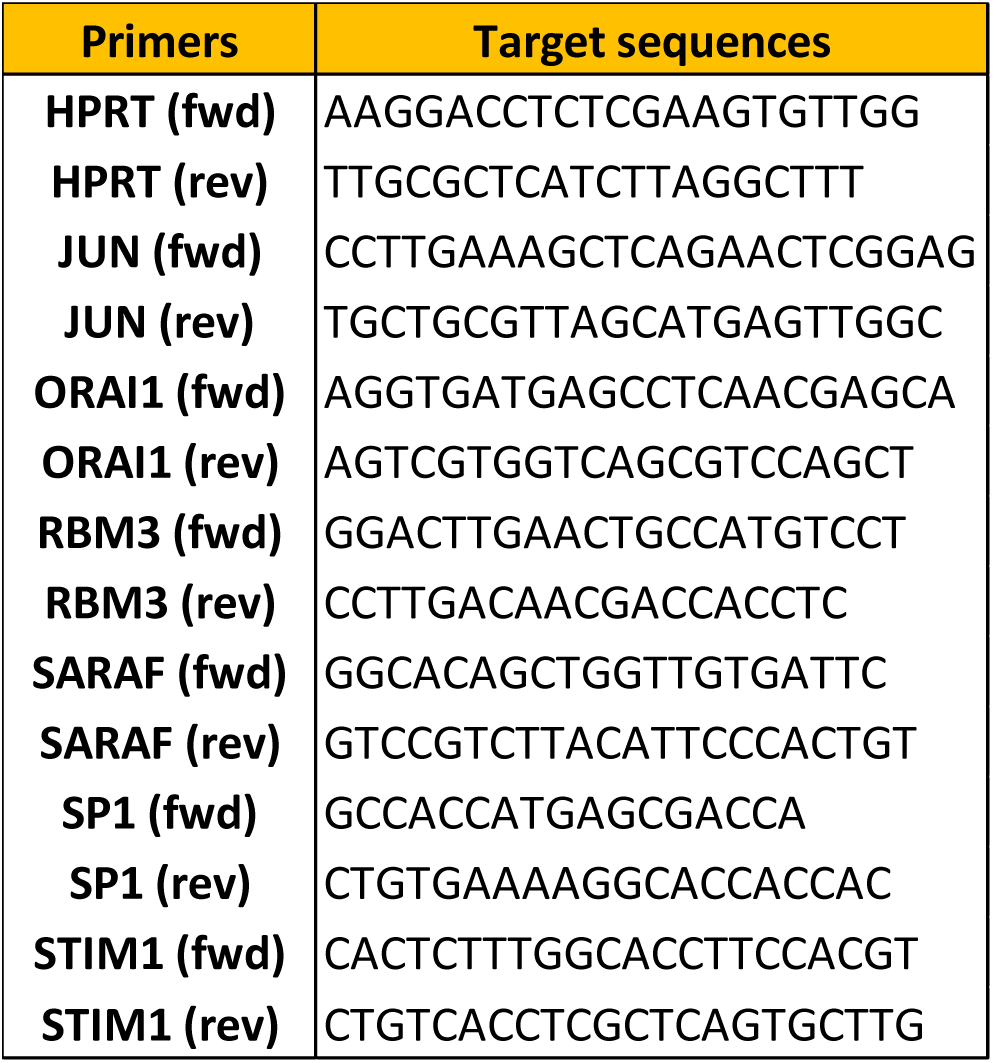
RT-qPCR primers used for validation of endogenous loci.

## Supplementary Figures

**Supplementary Figure 1.**
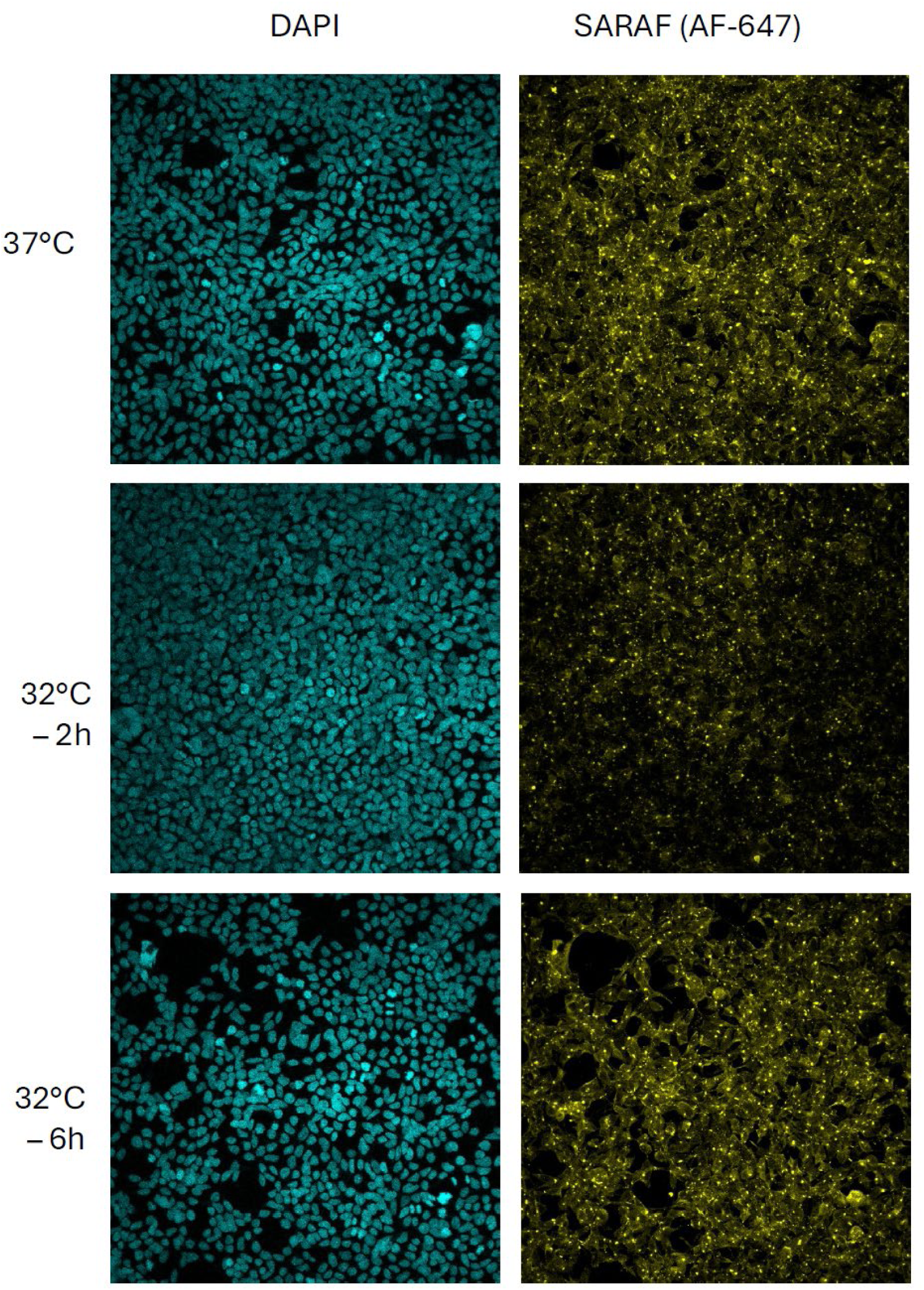
Changes in localization of SARAF upon a hypothermia stimulus. Representative images showing changes in distribution of the SARAF signal at 37 °C (top) and after a 2 hour (middle) or a six hour (bottom) exposure time to hypothermia (32 °C). Quantification of a multitude of such images is provided in **Figure 2C**.

**Supplementary Figure 2.**
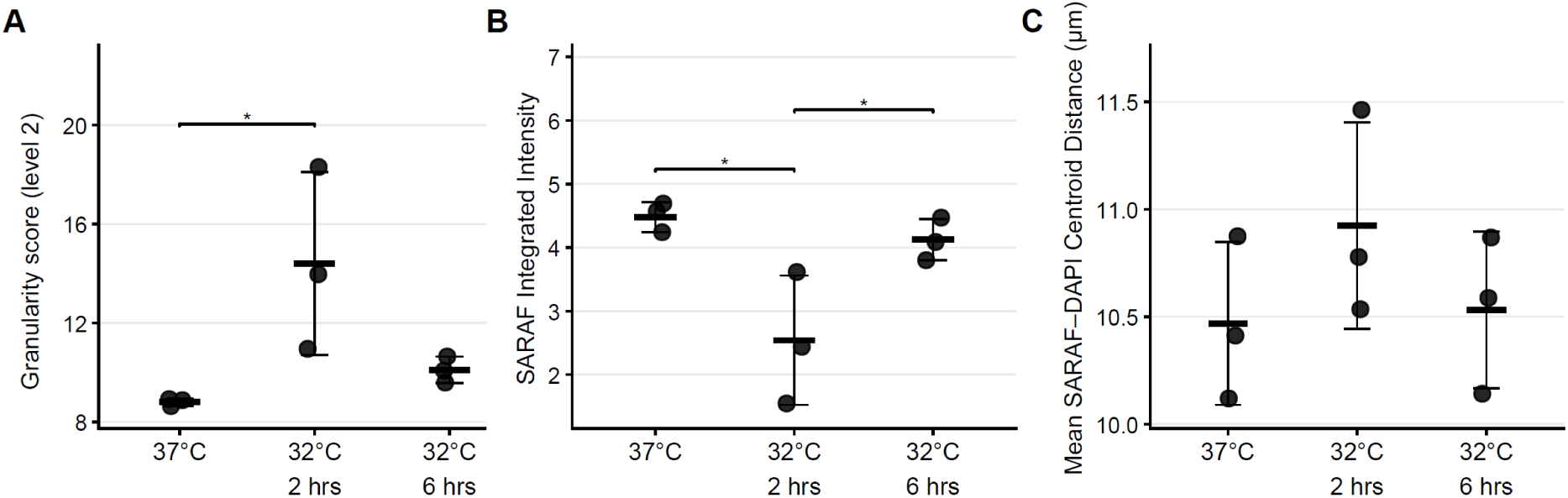
Granularity analysis revealed a redistribution of SARAF signal. (A) Quantitative analysis of SARAF fluorescence distribution across discrete granularity feature dimensions (ranging from low-order Level 2 to high-order Level 12) in HEK293 cells under normothermia and following specific duration of mild hypothermic exposure (32°C for 2 and 6 hours). Feature extraction was performed using a mathematical pattern-matching algorithm to segment localized signal variations. (B) Automated image-based quantification of total endogenous SARAF fluorescence intensity per cell across the indicated normothermic and hypothermic time points. (C) Subcellular spatial positioning analysis mapping the mean radial distance of cytoplasmic SARAF fluorescence intensity profiles relative to the nuclear boundary, as defined by DAPI. Data are displayed as mean ± SD (n= 3). Statistical significance was determined by one-way ANOVA followed by Tukey’s multiple comparisons test. \**p* < 0.05.

**Supplementary Figure 3.**
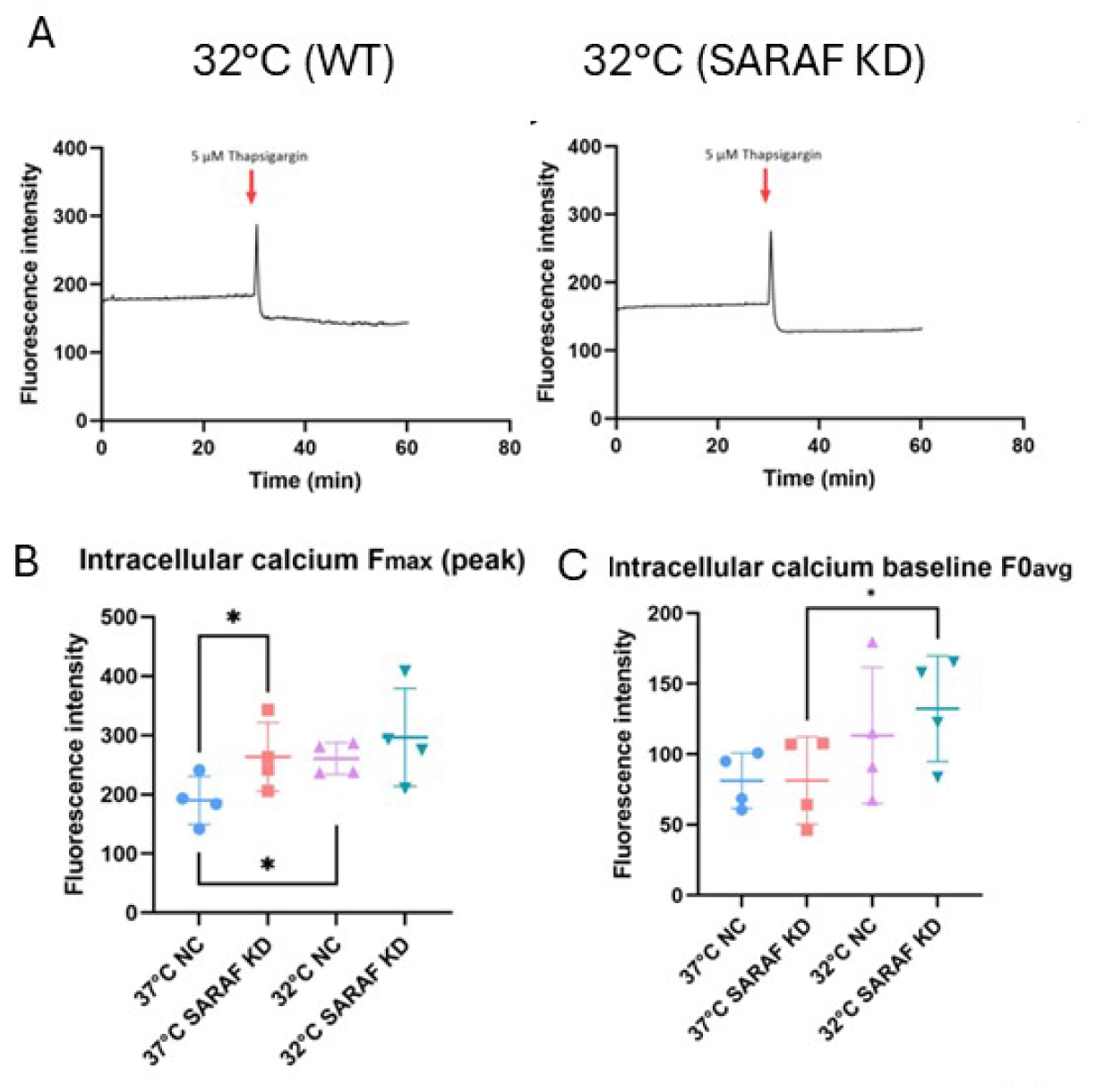
No significant changes upon SARAF KD in intracellular calcium dynamics during mild hypothermia. A) Quantification of intracellular calcium levels with TG exposure during hypothermia with (right) or without (left) *SARAF* KD. B) Intracellular calcium levels based on peak levels. C) Intracellular calcium levels based on baseline levels (prior to TG addition). Data are displayed as mean ± SD (n= 4). Statistical significance was evaluated by a Student’s t-test. Asterisks directly above the data points indicate comparisons with the controls (NC, 37 °C), while brackets delineate statistically significant differences between specified experimental groups. * indicates *p* < 0.05.

**Supplementary Figure 4.**
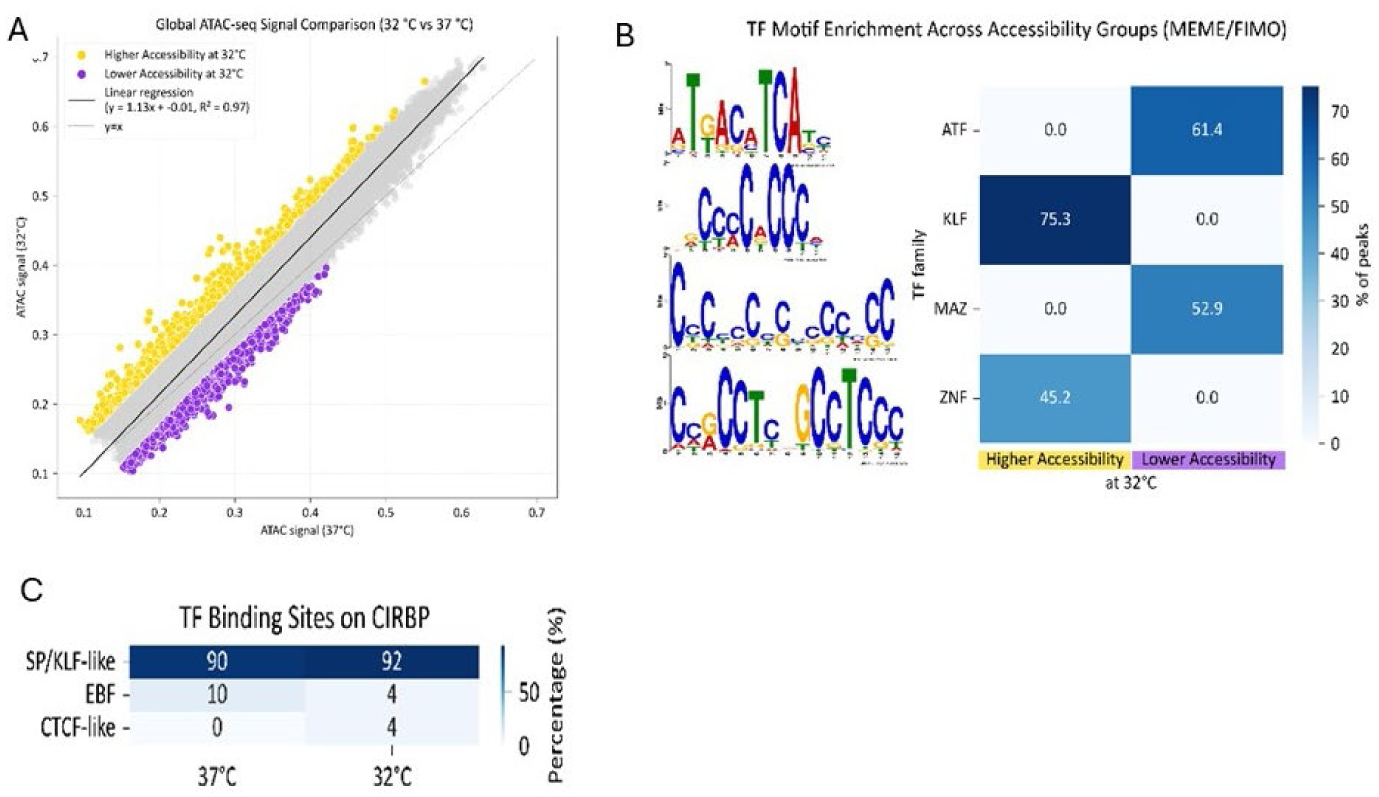
Chromatin accessibility and TF motif analysis under mild hypothermia. A) A global comparison of ATAC-seq signal between 32 °C and 37 °C across all peaks. Each point represents an ATAC-seq peak. A linear regression was fitted to model the global relationship between conditions (black line), with the diagonal (y = x) shown for reference. Peaks deviating from the regression line were classified based on residuals (±2.5 standard deviations) and an absolute accessibility difference threshold (|Δ accessibility| > 0.025). Peaks with higher-than-expected accessibility at 32 °C are shown in yellow, and those with lower-than-expected accessibility are shown in purple. B) TF motif enrichment across accessibility groups identified in A, determined using MEME/FIMO. Values represent the percentage of peaks containing motifs from each TF family in regions with increased or decreased accessibility at 32 °C. C) TF motif composition at CIRBP, another MHR factor known to be regulated at the transcriptional level.

**Supplementary Figure 5.**
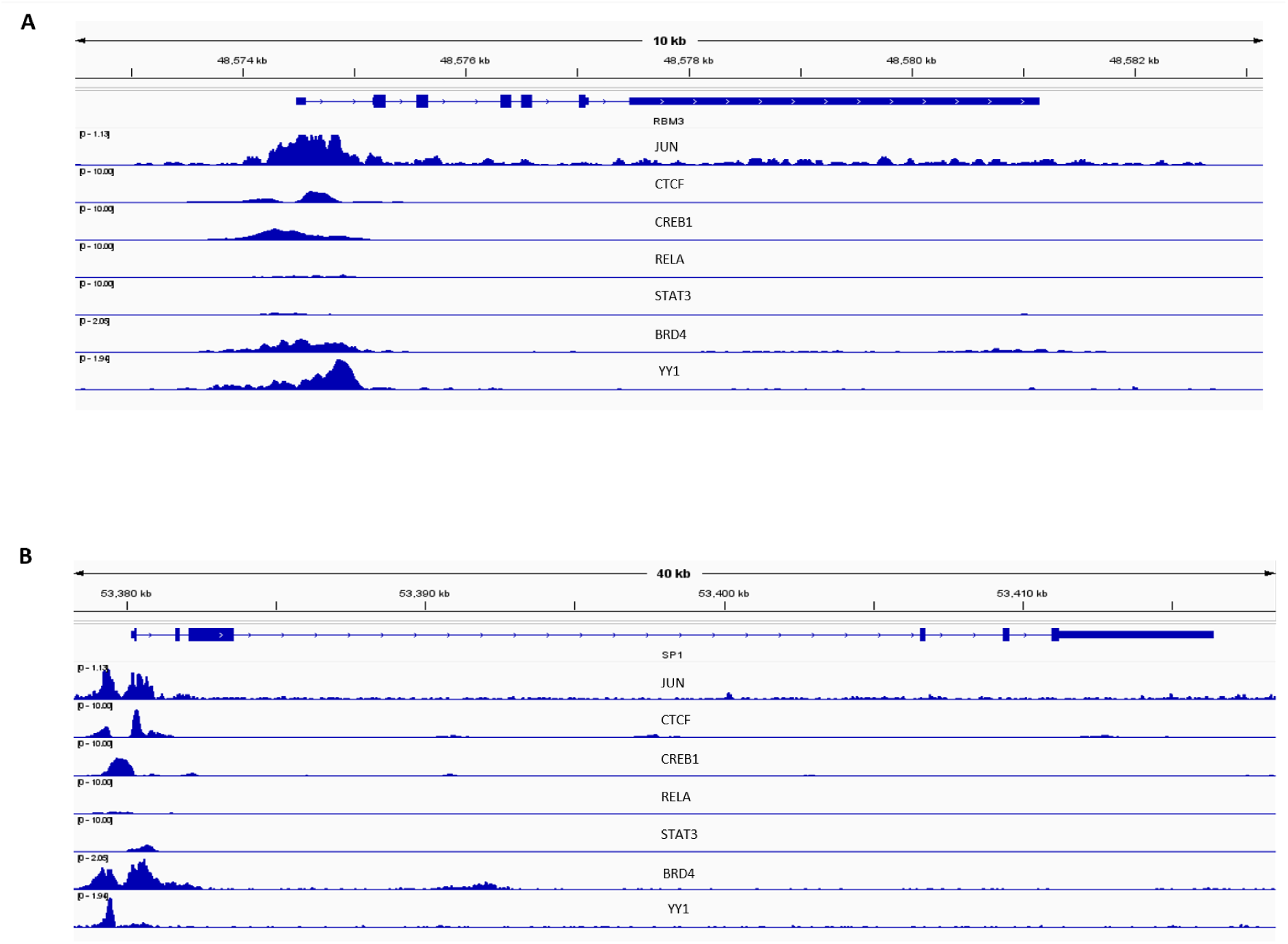
Shared TF binding at the promoter regions of *RBM3* and *SP1* (ChIP-Atlas). ChIP-Atlas binding peaks of the calcium-responsive TFs JUN, CTCF, CREB1, RELA, STAT3, BRD4 and YY1 at the promoter regions of *RBM3* (A) and *SP1* (B). Peak intensities reflect enrichment, with a threshold of 500 (−10 × log_10_ Q-value) applied to identify high-confidence binding events.

**Supplementary Figure 6.**
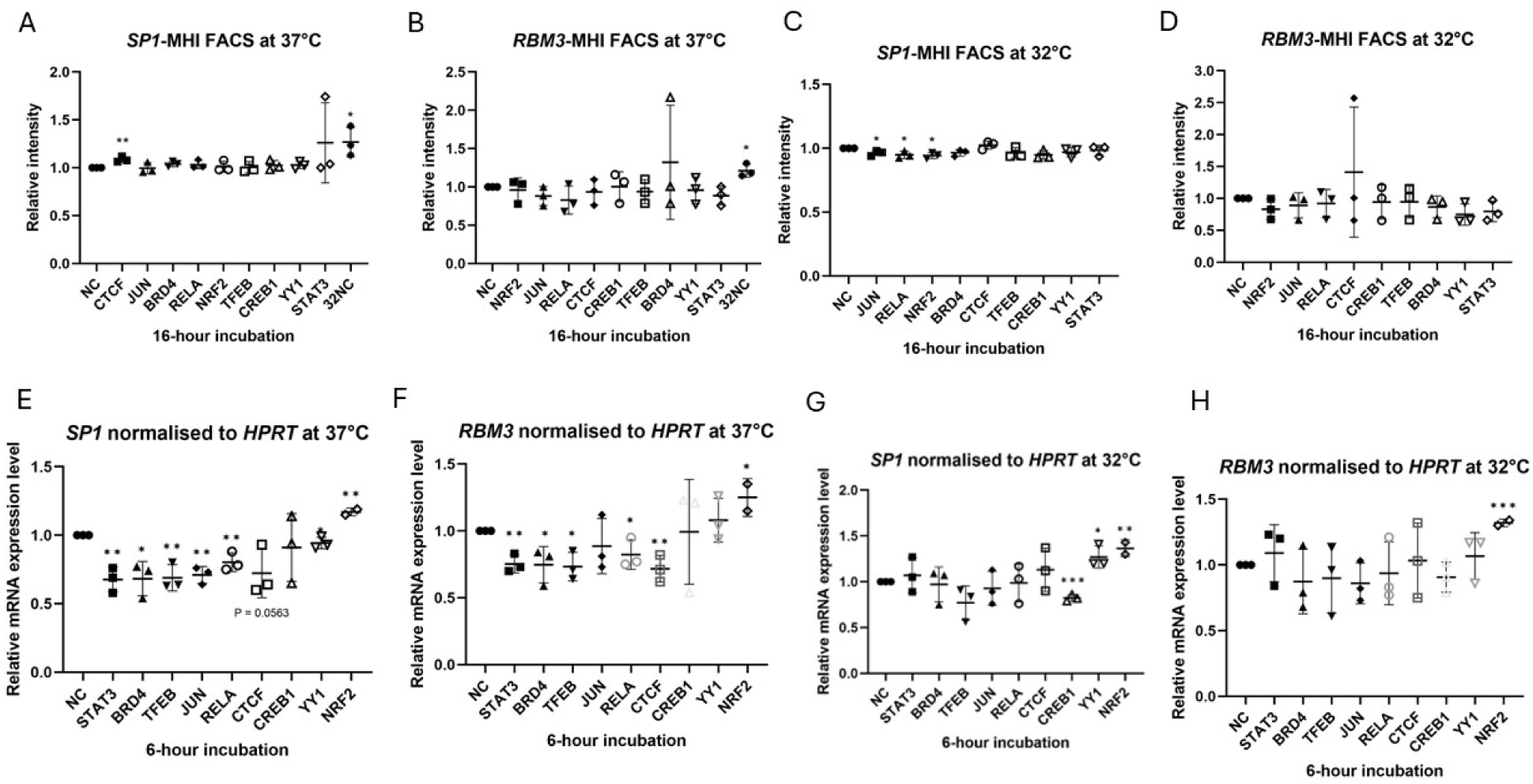
Validation of candidate TFs via MHI fluorescence and endogenous gene expression. (A-D) Impact of TF silencing on MHI activity. Flow cytometric analysis of (A) *SP1*-MHI and (B) *RBM3*-MHI fluorescence upon individual KD of 9 candidate TFs under normothermia. Corresponding analysis of (C) *SP1*-MHI and (D) *RBM3*-MHI fluorescence upon individual KD of 9 candidate TFs following 16 hours of mild hypothermic exposure. (E-H) Quantitative assessment of endogenous mRNA levels by RT-qPCR assays. Transcript abundance of (E) *SP1* and (F) *RBM3* following individual KD of 9 candidate TFs during normothermia. Parallel measurement of gene expression levels of (G) *SP1* and (H) *RBM3* after individual KD of 9 candidate TFs under mild hypothermic conditions. Data are presented as relative expression normalized to non-targeting control (NTC) siRNA. Data are displayed as mean ± SD (n= 3). Statistical significance was evaluated by a Student’s t-test. Asterisks directly above the data points indicate comparisons with the controls (NC, 37 °C). *, **, and *** denote *p* < 0.05, *p* < 0.01, and *p* < 0.001.

**Supplementary Figure 7.**
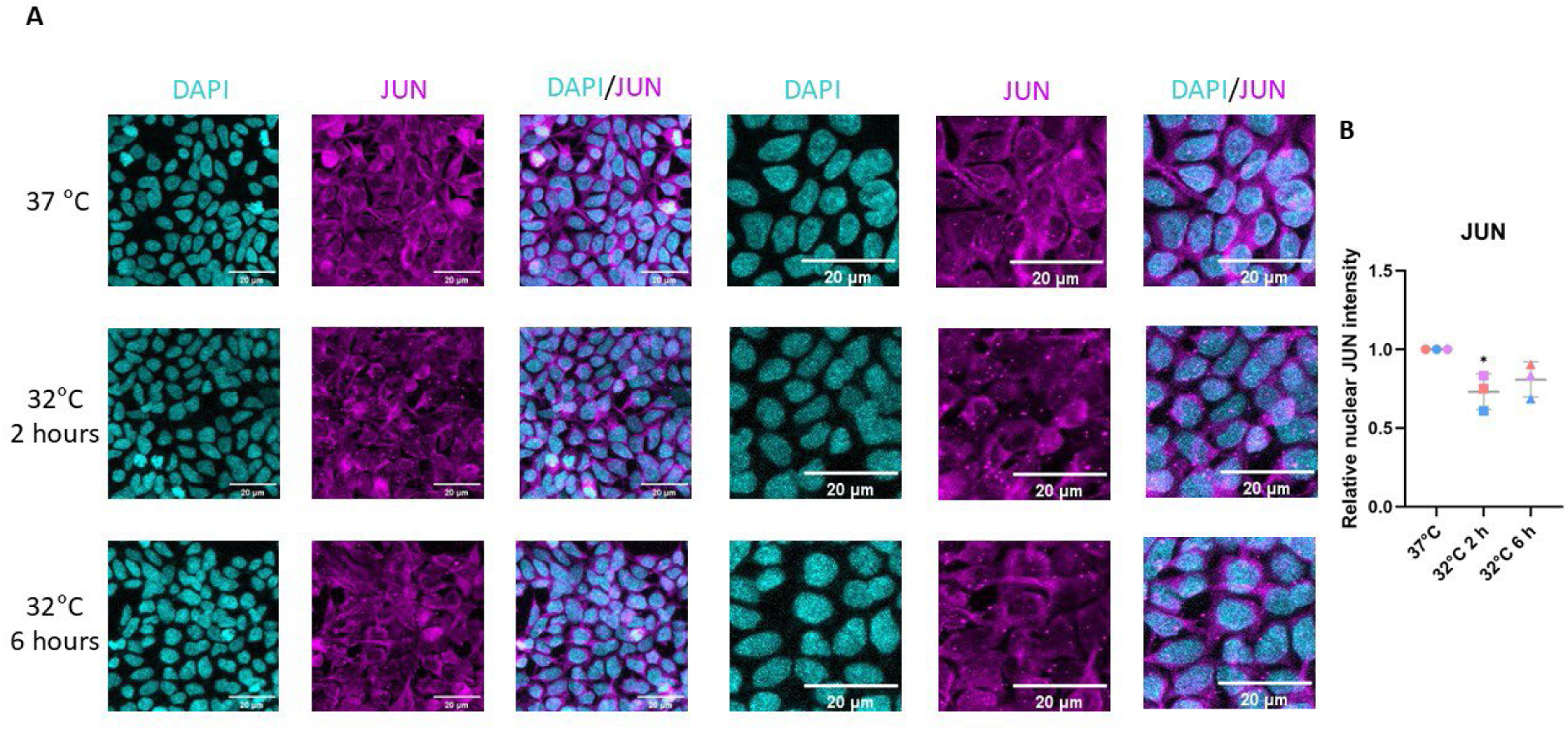
Nuclear JUN expression shows transient reduction upon hypothermia. (A-B) IF staining analysis of nuclear JUN expression in HEK293 cells subjected to time-course mild hypothermic conditions (0, 2, and 6 hours). (A) Representative images of JUN (magenta) and DAPI (blue) staining. (B) Quantification of nuclear JUN expression in at least 4-5 images per replicate in three biological replicates from panel A. Data are displayed as mean ± SD (n= 3). Statistical significance was determined by one-way ANOVA with Holm-Šídák’s multiple comparisons test for B. The asterisk directly above the data points indicates comparisons with the controls (37 °C). * indicates *p* < 0.05.

## Acknowledgements

The schematic image was designed by Servier Medical Art (https://smart.servier.com).

## Author contributions

H.T.B. and K.J. conceptualized and designed the study. H.T.B. received all the funding for this project. K.J., K.O., V.S., and K.J.A. performed all the experiments. K.J., K.J.A., K.O., and M.B. analyzed data. H.T.B. and K.J. interpreted the data. H.T.B. wrote the initial draft, K.J. revised the manuscript, and all authors reviewed, edited, and approved the final version of the manuscript.

## References

1. Protsiv, M., et al., Decreasing human body temperature in the United States since the industrial revolution. Elife, 2020. 9.

2. Mathew, J.L., N. Kaur, and J.M. Dsouza, Therapeutic hypothermia in neonatal hypoxic encephalopathy: A systematic review and meta-analysis. J Glob Health, 2022. 12: p. 04030.

3. Luedke, M.W., et al., Association of time-temperature curves with outcomes in temperature management for cardiac arrest. BMJ Neurol Open, 2022. 4(1): p. e000273.

4. Sun, Y.J., et al., Neuroprotection by Therapeutic Hypothermia. Front Neurosci, 2019. 13: p. 586.

5. Geurts, M., et al., COOLIST (Cooling for Ischemic Stroke Trial): A Multicenter, Open, Randomized, Phase II, Clinical Trial. Stroke, 2017. 48(1): p. 219–221.

6. Martini, W.Z., Coagulopathy by hypothermia and acidosis: mechanisms of thrombin generation and fibrinogen availability. J Trauma, 2009. 67(1): p. 202–8; discussion 208-9.

7. Wolberg, A.S., et al., A systematic evaluation of the effect of temperature on coagulation enzyme activity and platelet function. J Trauma, 2004. 56(6): p. 1221–8.

8. Fujita, J., Cold shock response in mammalian cells. J Mol Microbiol Biotechnol, 1999. 1(2): p. 243–55.

9. Knight, J.R., et al., Eukaryotic elongation factor 2 kinase regulates the cold stress response by slowing translation elongation. Biochem J, 2015. 465(2): p. 227–38.

10. Nagai, G. and Y. Nakaoka, Cooling sensitive [Ca2+]i response associated with signaling of G protein-coupled receptors. Biochem Biophys Res Commun, 1998. 248(3): p. 733–7.

11. Zhu, X., C. Bührer, and S. Wellmann, Cold-inducible proteins CIRP and RBM3, a unique couple with activities far beyond the cold. Cell Mol Life Sci, 2016. 73(20): p. 3839–59.

12. Bastide, A., et al., RTN3 Is a Novel Cold-Induced Protein and Mediates Neuroprotective Effects of RBM3. Curr Biol, 2017. 27(5): p. 638–650.

13. Preußner, M., et al., ASO targeting RBM3 temperature-controlled poison exon splicing prevents neurodegeneration in vivo. EMBO Mol Med, 2023. 15(5): p. e17157.

14. Yang, H.J., et al., Cold-inducible protein RBM3 mediates hypothermic neuroprotection against neurotoxin rotenone via inhibition on MAPK signalling. J Cell Mol Med, 2019. 23(10): p. 7010–7020.

15. Chip, S., et al., The RNA-binding protein RBM3 is involved in hypothermia induced neuroprotection. Neurobiol Dis, 2011. 43(2): p. 388–96.

16. Wierstra, I., Sp1: emerging roles--beyond constitutive activation of TATA-less housekeeping genes. Biochem Biophys Res Commun, 2008. 372(1): p. 1–13.

17. Sumitomo, Y., et al., Identification of a novel enhancer that binds Sp1 and contributes to induction of cold-inducible RNA-binding protein (cirp) expression in mammalian cells. BMC Biotechnol, 2012. 12: p. 72.

18. Rafnsdottir, S., et al., SMYD5 is a regulator of the mild hypothermia response. Cell Rep, 2024. 43(8): p. 114554.

19. Park, J., et al., SMYD5 methylation of rpL40 links ribosomal output to gastric cancer. Nature, 2024. 632(8025): p. 656–663.

20. Miao, B., et al., SMYD5 is a ribosomal methyltransferase that catalyzes RPL40 lysine methylation to enhance translation output and promote hepatocellular carcinoma. Cell Res, 2024. 34(9): p. 648–660.

21. Hamey, J.J., et al., SMYD5 is a ribosomal methyltransferase that trimethylates RPL40 lysine 22 through recognition of a KXY motif. Cell Rep, 2025. 44(4): p. 115518

22. Li, J., et al., Akt1-mediated CPR cooling protection targets regulators of metabolism, inflammation and contractile function in mouse cardiac arrest. PLoS One, 2019. 14(8): p. e0220604.

23. Yan, C., et al., Neuroprotective effects of mild hypothermia against traumatic brain injury by the involvement of the Nrf2/ARE pathway. Brain Behav, 2022. 12(8): p. e2686.

24. Fernandez-Marcos, P.J. and J. Auwerx, Regulation of PGC-1á, a nodal regulator of mitochondrial biogenesis. Am J Clin Nutr, 2011. 93(4): p. 884s–90.

25. Palty, R., et al., SARAF inactivates the store operated calcium entry machinery to prevent excess calcium refilling. Cell, 2012. 149(2): p. 425–38.

26. Zhou, Y., et al., STIM1 gates the store-operated calcium channel ORAI1 in vitro. Nat Struct Mol Biol, 2010. 17(1): p. 112–6.

27. Beer, M.A., D. Shigaki, and D. Huangfu, Enhancer Predictions and Genome-Wide Regulatory Circuits. Annu Rev Genomics Hum Genet, 2020. 21: p. 37–54.

28. Bailey TL, Machanick P. Inferring direct DNA binding from ChIP-seq. Nucleic Acids Res. 2012. 40(17):e128.

29. Kaczynski J, Cook T, Urrutia R. Sp1- and Krüppel-like transcription factors. Genome Biol. 2003. 4(2):206.

30. Karin M, Liu Zg, Zandi E. AP-1 function and regulation. Curr Opin Cell Biol. 1997. 9(2):240–6.

31. Nicolás, M., et al., Cloning and characterization of the 5’-flanking region of the human transcription factor Sp1 gene. J Biol Chem, 2001. 276(25): p. 22126–32.

32. Medina, D.L., et al., Lysosomal calcium signalling regulates autophagy through calcineurin and TFEB. Nat Cell Biol, 2015. 17(3): p. 288–99.

33. Bruna, B., et al., The signaling pathways underlying BDNF-induced Nrf2 hippocampal nuclear translocation involve ROS, RyR-Mediated Ca(2+) signals, ERK and PI3K. Biochem Biophys Res Commun, 2018. 505(1): p. 201–207.

34. Zou, Z., T. Ohta, and S. Oki, ChIP-Atlas 3.0: a data-mining suite to explore chromosome architecture together with large-scale regulome data. Nucleic Acids Res, 2024. 52(W1): p. W45–w53.

35. Filtz, T.M., W.K. Vogel, and M. Leid, Regulation of transcription factor activity by interconnected post-translational modifications. Trends Pharmacol Sci, 2014. 35(2): p. 76–85.

36. Lamph WW, Wamsley P, Sassone-Corsi P, Verma IM. Induction of proto-oncogene JUN/AP-1 by serum and TPA. Nature. 1988. 334(6183):629–31.

37. Si, W., et al., RNA Binding Protein Motif 3 Inhibits Oxygen-Glucose Deprivation/Reoxygenation-Induced Apoptosis Through Promoting Stress Granules Formation in PC12 Cells and Rat Primary Cortical Neurons. Front Cell Neurosci, 2020. 14: p. 559384.

38. Wu, L., et al., Therapeutic Hypothermia Enhances Cold-Inducible RNA-Binding Protein Expression and Inhibits Mitochondrial Apoptosis in a Rat Model of Cardiac Arrest. Mol Neurobiol, 2017. 54(4): p. 2697–2705.

39. Martina, J.A., et al., TFEB and TFE3 are novel components of the integrated stress response. Embo j, 2016. 35(5): p. 479–95.

40. Rao, A., Signaling to gene expression: calcium, calcineurin and NFAT. Nat Immunol, 2009. 10(1): p. 3–5.

41. Albarran, L., et al., Dynamic interaction of SARAF with STIM1 and Orai1 to modulate store-operated calcium entry. Sci Rep, 2016. 6: p. 24452.

42. Xiao, B., et al., Temperature-dependent STIM1 activation induces Ca²+ influx and modulates gene expression. Nat Chem Biol, 2011. 7(6): p. 351–8.

43. Ma, J., et al., Calhm2 governs astrocytic ATP releasing in the development of depression-like behaviors. Mol Psychiatry, 2018. 23(4): p. 883–891.

44. Ryu, H., et al., Sp1 and Sp3 are oxidative stress-inducible, antideath transcription factors in cortical neurons. J Neurosci, 2003. 23(9): p. 3597–606.

45. Andrews, S., FastQC: A Quality Control Tool for High Throughput Sequence Data. Babraham Bioinformatics, Babraham Institute, Cambridge, UK, 2010.

46. Langmead, B. and S.L. Salzberg, Fast gapped-read alignment with Bowtie 2. Nat Methods, 2012. 9(4): p. 357–359.

47. Zhang, Y., et al., Model-based analysis of ChIP-Seq (MACS). Genome Biol, 2008. 9(9): p. R137.

48. Bailey TL, et al., The MEME Suite. Nucleic Acids Res. 2015, 43: W39–49.

49. Yu G, Wang L, He Q (2015). “ChIPseeker: an R/Bioconductor package for ChIP peak annotation, comparison and visualization.” Bioinformatics, 31(14), 2382–2383. DOI: 10.1093/bioinformatics/btv145

